# Aquaculture facility-specific microbiota shape the zebrafish gut microbiome

**DOI:** 10.1101/2025.09.04.674294

**Authors:** Kayla C. Evens, Ingrid Bakke, Brendan J.M. Bohannan

**Author notes:** Corresponding author: Evens. **Email addresses of authors:** Bakke; Bohannan.

## Abstract

**Background:** Environmental microbiomes, such as those in recirculating aquaculture systems (RAS), can play a key role in shaping host-associated microbial communities. In zebrafish (*Danio rerio*) research, these interactions can introduce uncontrolled sources of variation, potentially confounding experimental outcomes across multiple facilities. Despite widespread zebrafish use in microbiome studies, few have characterized the microbial composition of both tank water and fish across multiple independent facilities to evaluate the consequences of environmental microbiome variation on the host microbiome.

**Results:** We compared water and zebrafish gut microbiomes across five aquaculture facilities—two in the United States and three in Norway—using a nested sampling design and 16S rRNA gene sequencing. Alpha diversity was consistently higher in tank water than in fish guts, and beta diversity metrics revealed distinct clustering by sample type, facility, and location. Differences in microbial community composition were significant across facilities, with both water and fish samples exhibiting facility-specific profiles. Similarity Percentage analysis identified taxonomic groups driving these differences, while Fast Expectation-Maximization for Microbial Source Tracking detected measurable contributions of tank water microbiota to zebrafish gut communities. Bray-Curtis dissimilarity values were lowest between fish and water from the same tank and increased with geographic and facility distance, indicating local microbial overlap. Relative abundance patterns and ordination plots further supported distinct and structured microbial assemblages across systems.

**Conclusions:** This study demonstrates that zebrafish aquaculture systems harbor unique microbial communities shaped by both environmental and geographic factors, with tank water acting as a potential source of gut-associated microbes. These findings underscore the importance of incorporating environmental microbiome assessments into zebrafish experimental design, particularly for studies focused on host-microbe interactions. Without such consideration, unaccounted variation in environmental microbiota may affect microbiome composition and reduce cross-study reproducibility. Moving forward, standardized reporting of environmental conditions and microbial composition across facilities will be critical for strengthening reproducibility and interpretation in zebrafish microbiome research.

## Background

Research over the past several decades has highlighted the substantial influence of the microbiome on vertebrate host development and fitness, through mediation of nutrition, metabolism, physiology, and immunology, among others [1, 2]. However, significant variation in the microbiome can be observed both between and within hosts [3, 4]. To better understand why this variability may occur and how it impacts animal host health and function, it is critical to first determine how the microbiome is acquired and maintained.

The structure and composition of the host-microbiome are partially influenced by factors outside of the individual host, which include, but are not limited to, food, interactions with other hosts, and the environmental microbiome [3]. While factors such as diet have been well-studied, the contribution of the environmental microbiome to host-microbiome variation remains unclear. Previous studies have identified correlations between environmental factors and host microbiome variability, but few have directly addressed the potential for the environmental acquisition of microbes by simultaneously assessing both host and environmental microbiomes [5–7].

Zebrafish (*Danio rerio*) are widely used as laboratory animal models for investigating questions related to host-environment interactions due, in part, to their rapid development, high reproductive rate, and ability to be derived germ-free (reared without microbes) [8, 9]. However, the generalizability of conclusions made from zebrafish experiments depends on the reproducibility of results. Studies in laboratory animals, including mice [10, 11] and non-human primates [7, 12], have revealed significant microbiome variation associated with housing condition and vendor sources, even among genotype-matched individuals. If unaccounted for, such variation can confound cross-study comparisons, especially in microbiome-sensitive research.

Similar intra-facility [13] and inter-facility host-microbiome variation [14, 15] has been reported in zebrafish. However, to the best of our knowledge, no studies have directly examined the contribution of the environmental microbiome to zebrafish gut microbiome variability across multiple modern research aquaculture facilities.

Modern zebrafish aquaculture facilities typically use recirculating aquaculture systems (RAS), which, as the name implies, treat and reuse water within a closed-loop system. Over recent decades, interest in these systems has grown in both research and commercial settings, in part due to their efficiency and reduced environmental impact. Studies have demonstrated that these systems may maintain high populations of beneficial bacteria or, at the very least, reduce the abundance of rapid-growing heterotrophic bacteria potentially harmful to fish. This is related to long water retention times, the presence of a biofilter that consumes substrates for bacterial growth, and similar carrying capacities and bacterial densities throughout the system-resulting in selection for slow-growing but competitively superior non-pathogens [16–18]. However, technical specifications for water treatment can vary, with some facilities instead employing flow-through systems, in which all outflow water is discharged rather than recirculated.

Despite a general shared goal of maintaining clean, safe rearing conditions for zebrafish, differences in external inputs and management practices are likely to contribute to variation in water microbial community composition across aquaculture facilities. As such, if zebrafish microbiomes are largely shaped by environmental acquisition, facility-level differences in water microbial communities could drive significant variation in the zebrafish gut microbiome. However, research suggests that the zebrafish intestine is at least somewhat selective, with a “core” intestinal microbiome reported across different facilities [14, 15]. Therefore, selection by the zebrafish gut could mitigate the effect of environmental microbiome variability on the composition of the gut microbiome.

This study addresses two main questions: (1) How do the environmental and fish gut microbiomes vary across aquaculture facilities? and (2) Can we identify patterns of covariation between the environmental microbiome and the zebrafish gut microbiome? To investigate these questions, we selected five facilities in two geographic locations—Trondheim, Norway and Eugene, Oregon—that are representative of typical zebrafish research environments. Within each location, facilities are located within 300 meters of each other on the University of Oregon (UO) campus and within 700 meters on the Norwegian University of Science and Technology (NTNU) campus (Figure 1). These facilities predominantly utilize recirculating aquaculture systems (RAS), though one facility (Nor2B) employs a flow-through system with a different water treatment approach. We specifically focused on facilities that supply zebrafish for basic research, as these facilities tend to share operational similarities due to the sensitive nature of rearing laboratory animals, allowing us to better isolate the interaction between host and environmental microbiomes while providing insight into the reproducibility of results from zebrafish sourced from different locations.

**Figure 1:**
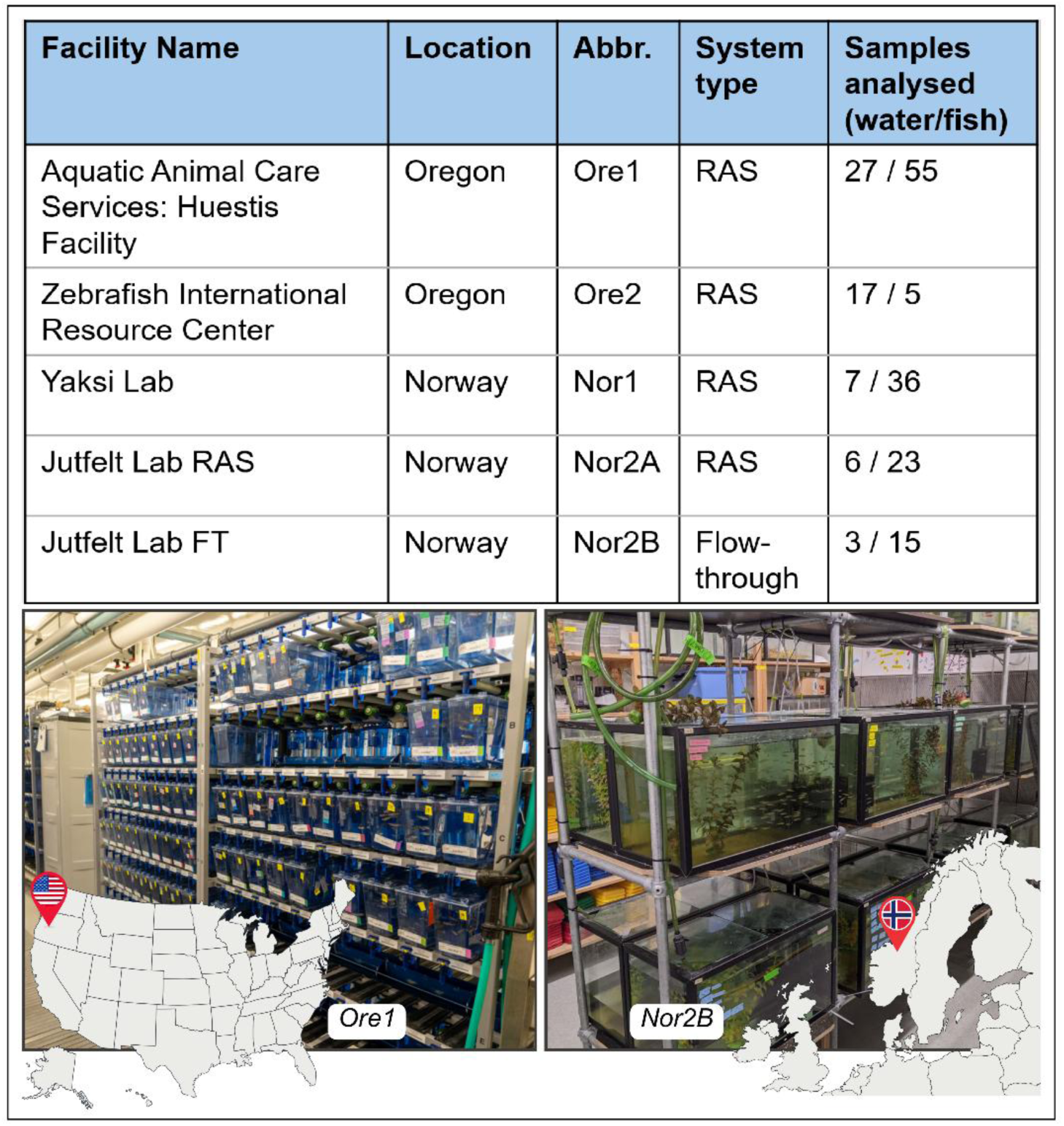
Aquaculture facilities (facility name, geographic location, abbreviation, and water system type) and samples collected from tank water and zebrafish. The two photos of zebrafish tanks (Ore1-left; Nor2B-right) provide an example of the variety and organization of tanks common in modern aquaculture facilities. *Image of Ore1 facility courtesy of Kelley Christensen, 2024, University of Oregon. Image of Nor2B facility, Kayla C. Evens, 2022*.

By using 16S rRNA gene-sequencing to quantify microbial variation in paired samples of fish and tank water from these five facilities, we demonstrate significant variation in both the water and fish gut microbiomes, despite the relative technical similarities of their water systems. Furthermore, we show that overall, the microbiomes of fish and water are more similar within facilities than between facilities. Finally, using source tracking methods, we demonstrate a consistent influence of the environmental microbiome on the zebrafish gut microbiome.

## Results

### Tank Water Microbiota

This study aimed to assess how variation in the tank water microbiome influences the zebrafish gut microbiome across aquaculture facilities. Before addressing this, we first needed to determine whether there was observable variation in the tank water microbiome across the five facilities. We analyzed metrics of both community composition and diversity within and across 50 tank water samples, identifying a total of 1,625 unique amplicon sequence variants (ASVs) after normalizing the data to 1,369 reads per sample (*n* = Ore1 (27); Ore2 (17); Nor1 (7); Nor2A (6); Nor2B (3)).

#### Variation in tank water microbiota is driven by shifts in dominant bacterial taxa

To compare water microbial diversity within facilities, we first examined α-diversity metrics. Shannon-Wiener index values did not differ significantly across facilities (Kruskal-Wallis: *p* = 0.13), though pairwise comparisons revealed significant differences between Nor2A and Nor1 (pairwise Wilcoxon test: *p* = 0.008) and between Nor2B and Nor1 (*p* = 0.017). However, the Inverse Simpson index, which emphasizes numerically dominant taxa, varied significantly across facilities (Kruskal-Wallis: *p* = 0.029). Pairwise tests showed significant differences between Nor1 and both Ore1 (pairwise Wilcoxon test: *p* = 0.004) and Ore2 (*p* = 0.013) (Figure 2). Overall, the Nor1 facility exhibited significantly higher α-diversity than the Nor2A and Nor2B facilities.

**Figure 2:**
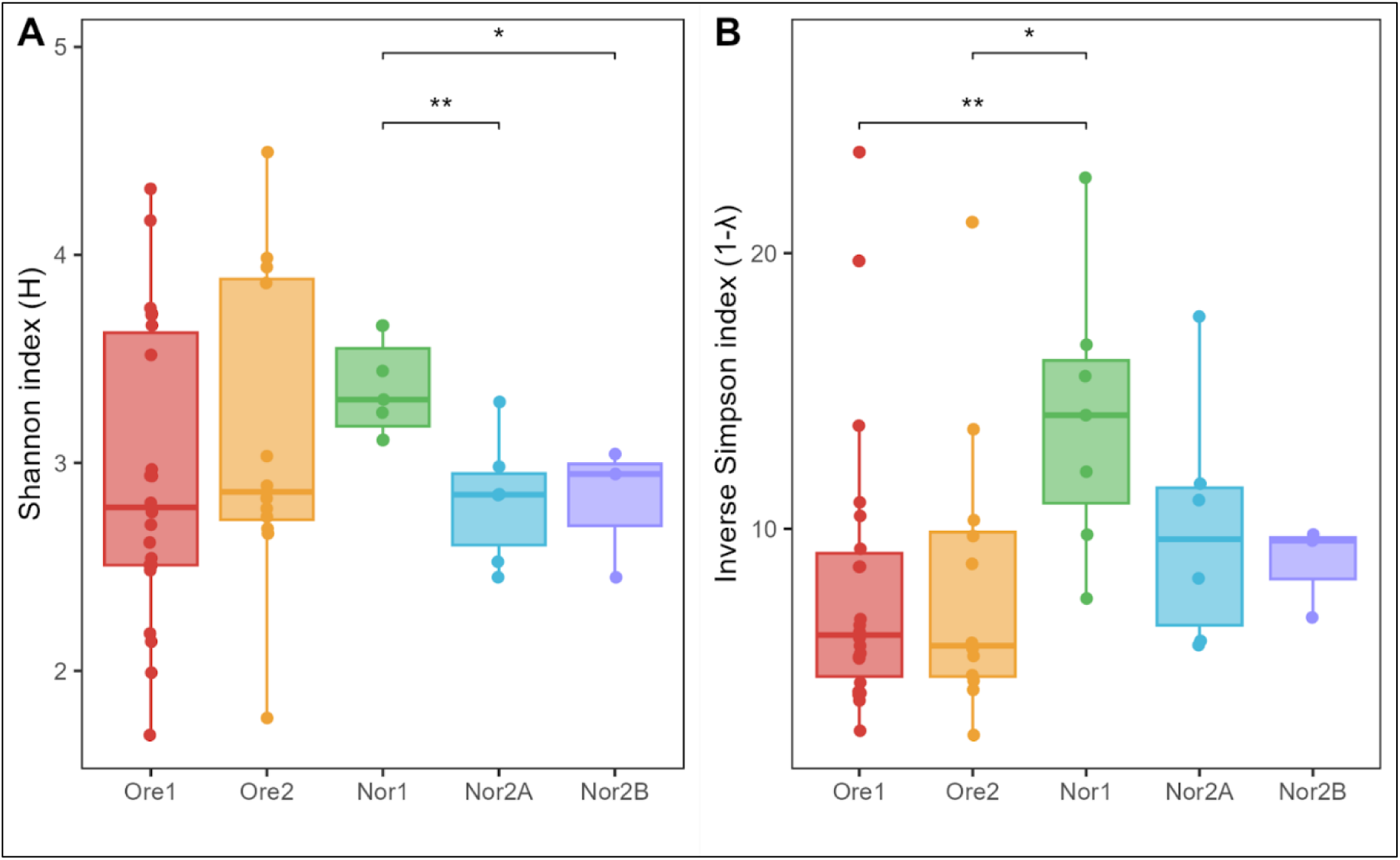
Alpha diversity metrics of tank water microbiota with (A) Shannon diversity and (B) Inverse Simpson diversity. Tukey-style box and whisker plots display the median (center horizontal line) and interquartile range (upper and lower bounds of the box), while whiskers extend +/- 1.5 times the interquartile range. Significance calculated with pairwise Wilcoxon rank-sum test with Benjamini-Hochberg p-value correction (* = p < 0.05, ** = p < 0.01, *** = p < 0.001, **** = p < 0.0001, ns = non-significant).

Given that differences appeared to be largely driven by dominant taxa, we further examined tank water microbial community composition across facilities. Though no single phylum was shared among all tank water samples, seven phyla were detected at a mean relative abundance threshold of 1%. Proteobacteria was the dominant phylum across all facilities, comprising between 52% (Ore1) and 92% (Nor2A) of the water microbiome. There were additionally 19 abundant (>1%) genera represented across all facilities. Of these abundant taxa, none were unique to water samples, as all were also present in zebrafish gut samples. The Ore1 and Ore2 facility water microbiomes were both dominated by *Cetobacterium* (20.2% and 42% of sequences, respectively). Ore1 was further characterized by high relative abundances of *Psychrobacter* (19%) and *Aeromonas* (12%). In contrast, *Pseudomonas* was dominant in Nor1 (35%) while *Rheinheimera* was most prevalent in Nor2A (33%). Compared to the other facilities, Nor2B exhibited no single dominant taxon, instead featuring multiple Proteobacteria, including *Acidovorax* (11%), *Nevskia* (10%), and *Limnobacter* (9%) among others (Figure 3).

**Figure 3:**
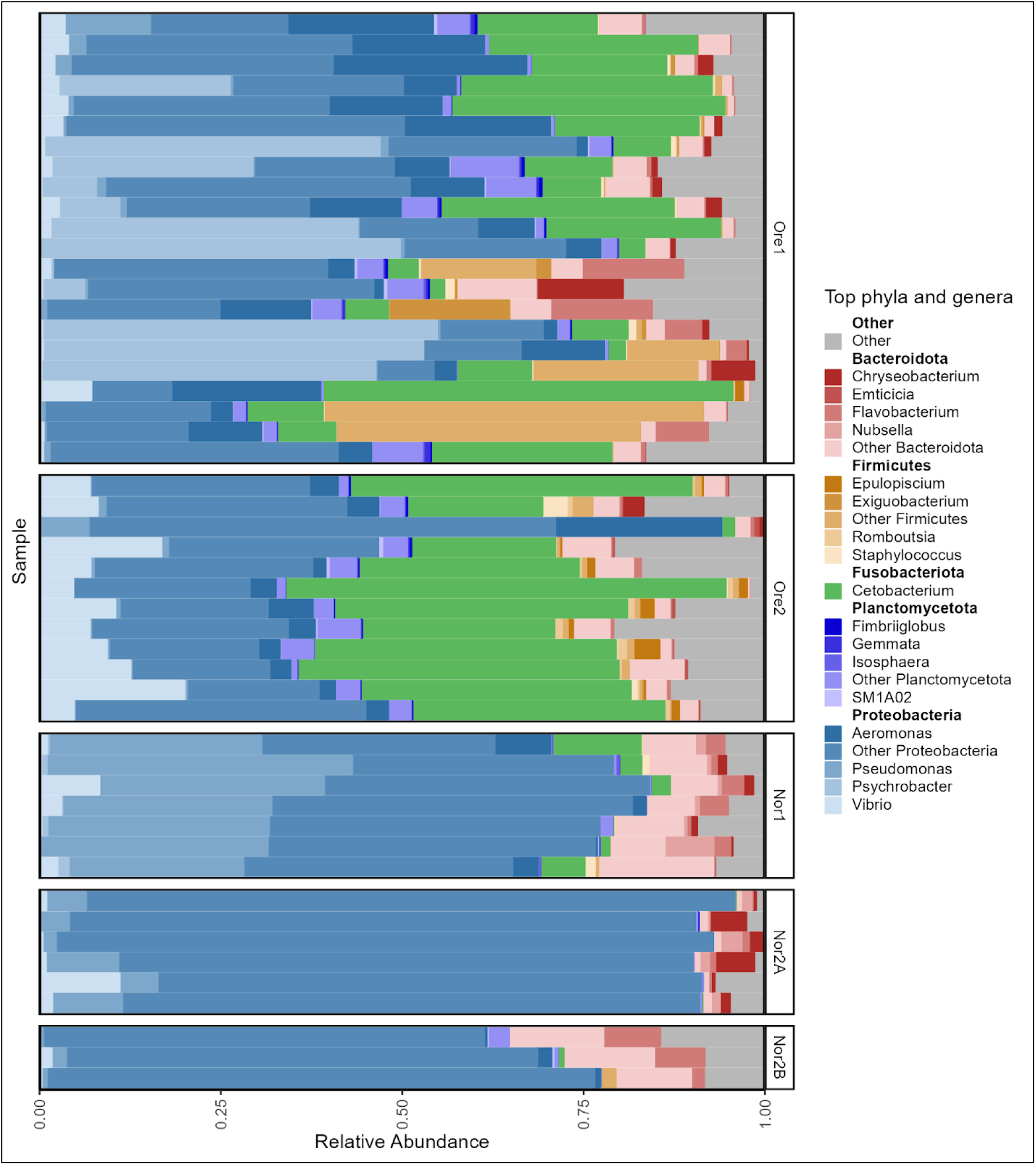
Phylum and genus-level taxonomic composition of tank water microbiome samples across aquaculture facilities. Each bar represents one tank water sample. The mean relative abundance is grouped by aquaculture facility and is displayed for only the top five most abundant phyla and top four genera within each phylum as averaged across samples.

#### Aquaculture facilities are a major driver of tank water microbial variation

Tank water microbiome composition varied significantly across facilities. Given the nested structure of our study design (facilities nested within geographic locations), we conducted permutational analysis of variance (PERMANOVA) using a hierarchical approach. The initial analysis testing the combined effects of location (Norway vs. Oregon), facility, and fish genotype status (held within the tank) revealed significant effects for all factors (Bray-Curtis and unweighted UniFrac: *p* < 0.002). Location explained 20.4% and 12.8% of variation in Bray-Curtis and unweighted UniFrac distances, respectively, while individual facility identity accounted for 22.7% and 18% of the variation.

To properly assess the effect of fish genotype status while accounting for the nested design, we performed a constrained PERMANOVA with permutations restricted within each facility. This analysis revealed that genotype status alone did not significantly influence water microbiome composition (Bray-Curtis: *p* = 0.567, unweighted UniFrac: *p* = 0.427), suggesting that the apparent genotype effect in the initial hierarchical model was confounded with facility-level differences rather than representing a true biological effect of fish genetics on the water microbial community (Supplementary Table 1).

Pairwise PERMANOVA comparisons revealed stronger differentiation between geographic locations than within locations. While all facility pairs showed significant differences (*p* < 0.05), Norwegian facilities exhibited weaker differentiation among themselves, particularly between Nor2A and Nor2B (*p*-adj. = 0.11) and between Nor2B and Nor1 (*p*-adj. = 0.10), suggesting greater similarity in water microbiome composition within the Norwegian location (Table 1).

**Table 1:**
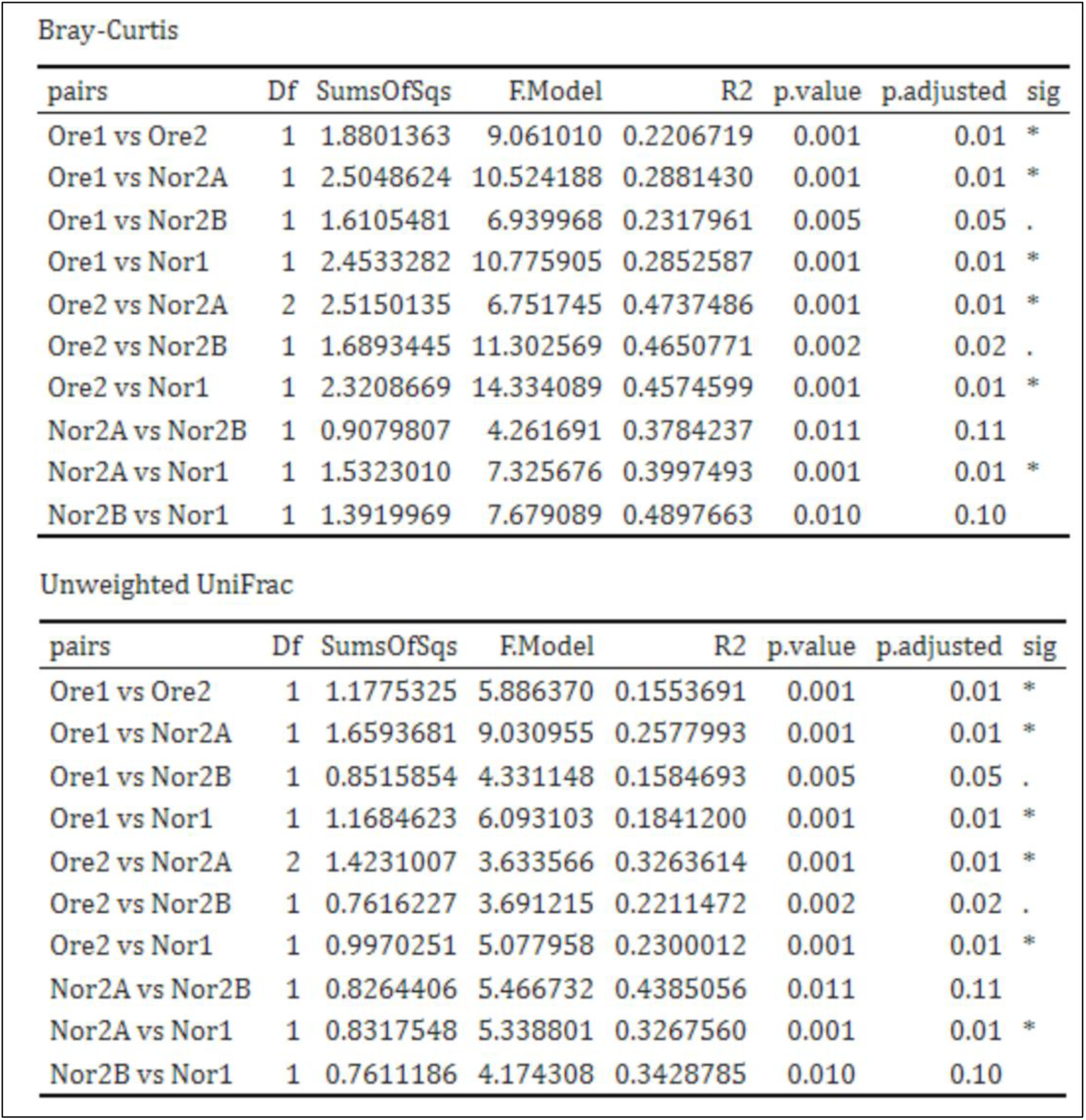
Pairwise PERMANOVA on Bray-Curtis and unweighted UniFrac distance of aquaculture facility water microbiome. p-value adjusted with Benhamini-Hochberg correction.

| Bray-Curtis |  |  |  |  |  |  |  |
| --- | --- | --- | --- | --- | --- | --- | --- |
| pairs | Df | SumsOfSqs | FModel | R2 | p.value | p.adjusted | sig |
| Ore1 vs Ore2 | 1 | 1.8801363 | 9.061010 | 0.2206719 | 0.001 | 0.01 | * |
| Ore1 vs Nor2A | 1 | 2.5048624 | 10.524188 | 0.2881430 | 0.001 | 0.01 | * |
| Ore1 vs Nor2B | 1 | 1.6105481 | 6.939968 | 0.2317961 | 0.005 | 0.05 | . |
| Ore1 vs Nor1 | 1 | 2.4533282 | 10.775905 | 0.2852587 | 0.001 | 0.01 | * |
| Ore2 vs Nor2A | 2 | 2.5150135 | 6.751745 | 0.4737486 | 0.001 | 0.01 | * |
| Ore2 vs Nor2B | 1 | 1.6893445 | 11.302569 | 0.4650771 | 0.002 | 0.02 | . |
| Ore2 vs Nor1 | 1 | 2.3208669 | 14.334089 | 0.4574599 | 0.001 | 0.01 | * |
| Nor2A vs Nor2B | 1 | 0.9079807 | 4.261691 | 0.3784237 | 0.011 | 0.11 |  |
| Nor2A vs Nor1 | 1 | 1.5323010 | 7.325676 | 0.3997493 | 0.001 | 0.01 | * |
| Nor2B vs Nor1 | 1 | 1.3919969 | 7.679089 | 0.4897663 | 0.010 | 0.10 |  |

| Unweighted UniFrac |  |  |  |  |  |  |  |
| --- | --- | --- | --- | --- | --- | --- | --- |
| pairs | Df | SumsOfSqs | FModel | R2 | p.value | p.adjusted | sig |
| Ore1 vs Ore2 | 1 | 1.1775325 | 5.886370 | 0.1553691 | 0.001 | 0.01 | * |
| Ore1 vs Nor2A | 1 | 1.6593681 | 9.030955 | 0.2577993 | 0.001 | 0.01 | * |
| Ore1 vs Nor2B | 1 | 0.8515854 | 4.331148 | 0.1584693 | 0.005 | 0.05 | . |
| Ore1 vs Nor1 | 1 | 1.1684623 | 6.093103 | 0.1841200 | 0.001 | 0.01 | * |
| Ore2 vs Nor2A | 2 | 1.4231007 | 3.633566 | 0.3263614 | 0.001 | 0.01 | * |
| Ore2 vs Nor2B | 1 | 0.7616227 | 3.691215 | 0.2211472 | 0.002 | 0.02 | . |
| Ore2 vs Nor1 | 1 | 0.9970251 | 5.077958 | 0.2300012 | 0.001 | 0.01 | * |
| Nor2A vs Nor2B | 1 | 0.8264406 | 5.466732 | 0.4385056 | 0.011 | 0.11 |  |
| Nor2A vs Nor1 | 1 | 0.8317548 | 5.338801 | 0.3267560 | 0.001 | 0.01 | * |
| Nor2B vs Nor1 | 1 | 0.7611186 | 4.174308 | 0.3428785 | 0.010 | 0.10 |  |

PCoA plots of Bray-Curtis and unweighted UniFrac distances further corroborated these results, with tank water microbiomes clustering strongly by facility, though some overlap was observed among Norwegian facilities (Figure 4).

**Figure 4:**
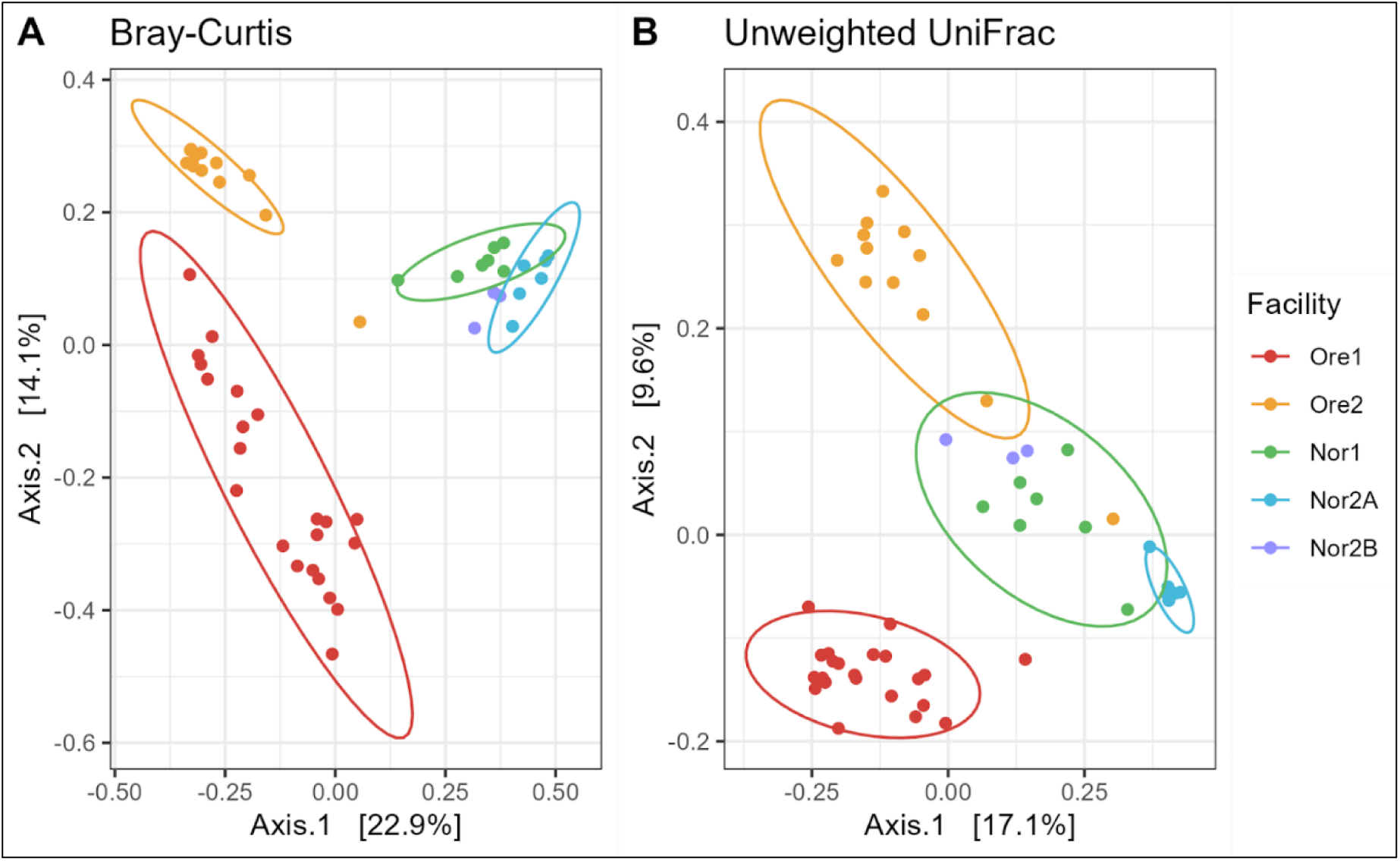
Beta diversity of tank water samples across aquaculture facilities. A principal coordinate analysis (PCoA) of (A) Bray-Curtis and (B) unweighted UniFrac dissimilarity. Colors represent facilities with normal confidence ellipses (95%).

To further investigate which genus-level taxa most contributed to differences observed between aquaculture facilities, we performed a Similarity Percentage (SIMPER) analysis [19].

The SIMPER analysis revealed that differences in tank water microbiota among facilities were primarily driven by shifts in members of the Fusobacteria and Proteobacteria phyla. *Cetobacterium* (Fusobacteria) was a major contributor to dissimilarity, accounting for 18-20% of the observed differences between Ore2 and all three Norwegian facilities (Nor1, Nor2A, Nor2B), and 13% of the differences between the two Oregon facilities (Ore1 vs. Ore2). Proteobacteria taxa also contributed substantially to facility differences, with *Pseudomonas* and *Rheinheimera* together responsible for 13-17% of the dissimilarity between Ore1 and the Norwegian facilities. These results indicate that both geographic and facility-specific factors influence water microbiome composition, with certain taxa consistently driving differences across multiple facility comparisons (Supplementary Table 2).

### Zebrafish Gut Microbiota

To determine whether zebrafish gut microbiomes varied across tanks and facilities, we applied similar analyses of community composition and diversity. Across 134 fish samples, we identified 1,281 unique ASVs after data were normalized to 1,171 reads per sample (*n* = Ore1 (55); Ore2 (5); Nor1 (15); Nor2A (36); Nor2B (23)).

#### Differences in zebrafish gut microbiota are driven by shifts in dominant bacterial taxa

Zebrafish gut microbiomes varied significantly in α-diversity metrics, both across all facilities (Kruskal-Wallis: Shannon: *p* < 0.001; Inverse Simpson: *p* = 0.003) and between pairs of facilities (pairwise Wilcoxon test: Shannon: Ore1 vs. Nor2A: *p* < 0.001; Ore2 vs. Nor2A, *p* < 0.001; Nor2A vs. Nor2B, *p* <0.001; Nor2A vs. Nor1, *p* < 0.001, Inverse Simpson: Ore1 vs. Nor2A: *p* = 0.008; Ore2 vs. Nor2A, *p* < 0.001; Nor2A vs. Nor2B, *p* = 0.002; Nor2A vs. Nor1, *p* < 0.001). Alpha diversity was consistently lowest in the Nor2A facility (Figure 5).

**Figure 5:**
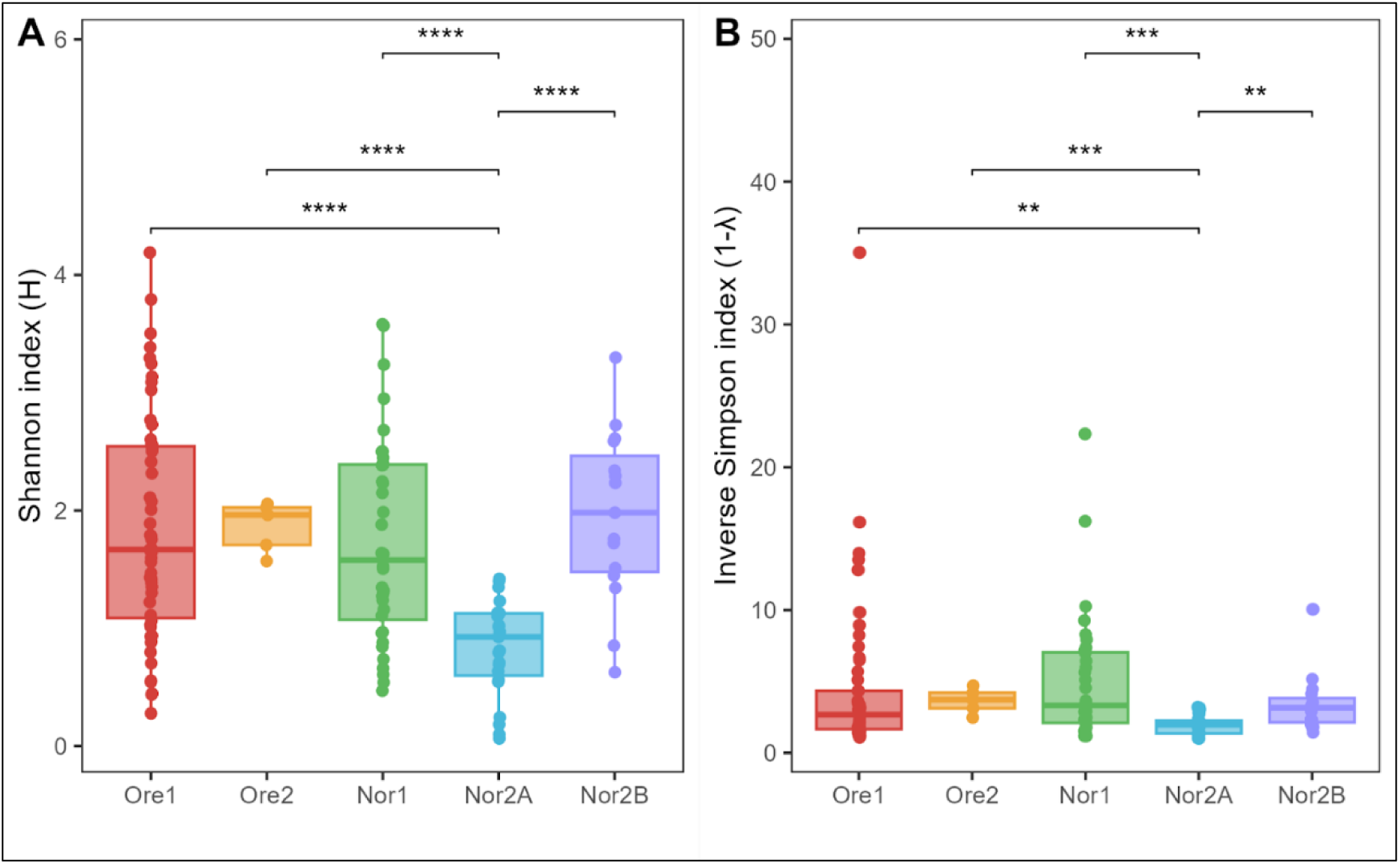
Alpha diversity metrics of zebrafish gut microbiota with (A) Shannon diversity and (B) Inverse Simpson diversity. Tukey-style box and whisker plots display the median (center horizontal line) and interquartile range (upper and lower bounds of the box), while whiskers extend +/- 1.5 times the interquartile range. Significance calculated with pairwise Wilcoxon rank-sum test with Benjamini-Hochberg p-value correction (* = p < 0.05, ** = p < 0.01, *** = p < 0.001, **** = p < 0.0001, ns = non-significant).

Fifteen genera exceeded a mean relative abundance threshold of 1% across all facilities, three of which (*Cetobacterium*, *Aeromonas*, and *Plesiomonas*) were shared among all sites. Additionally, of the identified genera, three were unique to the fish gut microbiome, all members of the order Lactobacillales (phylum Firmicutes)-*Lactococcus*, *Lactilactobacillus*, and *Pediococcus*.

In the Oregon facilities, fish gut microbiomes were dominated by *Cetobacterium (*56% in Ore1, 47% in Ore2), whereas *Aeromonas* was most abundant in Nor2B (60%) and Nor1(24%). The Nor2A fish gut microbiomes, in contrast, were dominated by *Vibrio* (47%), which was also abundant in all facilities except Nor1 (Figure 6).

**Figure 6:**
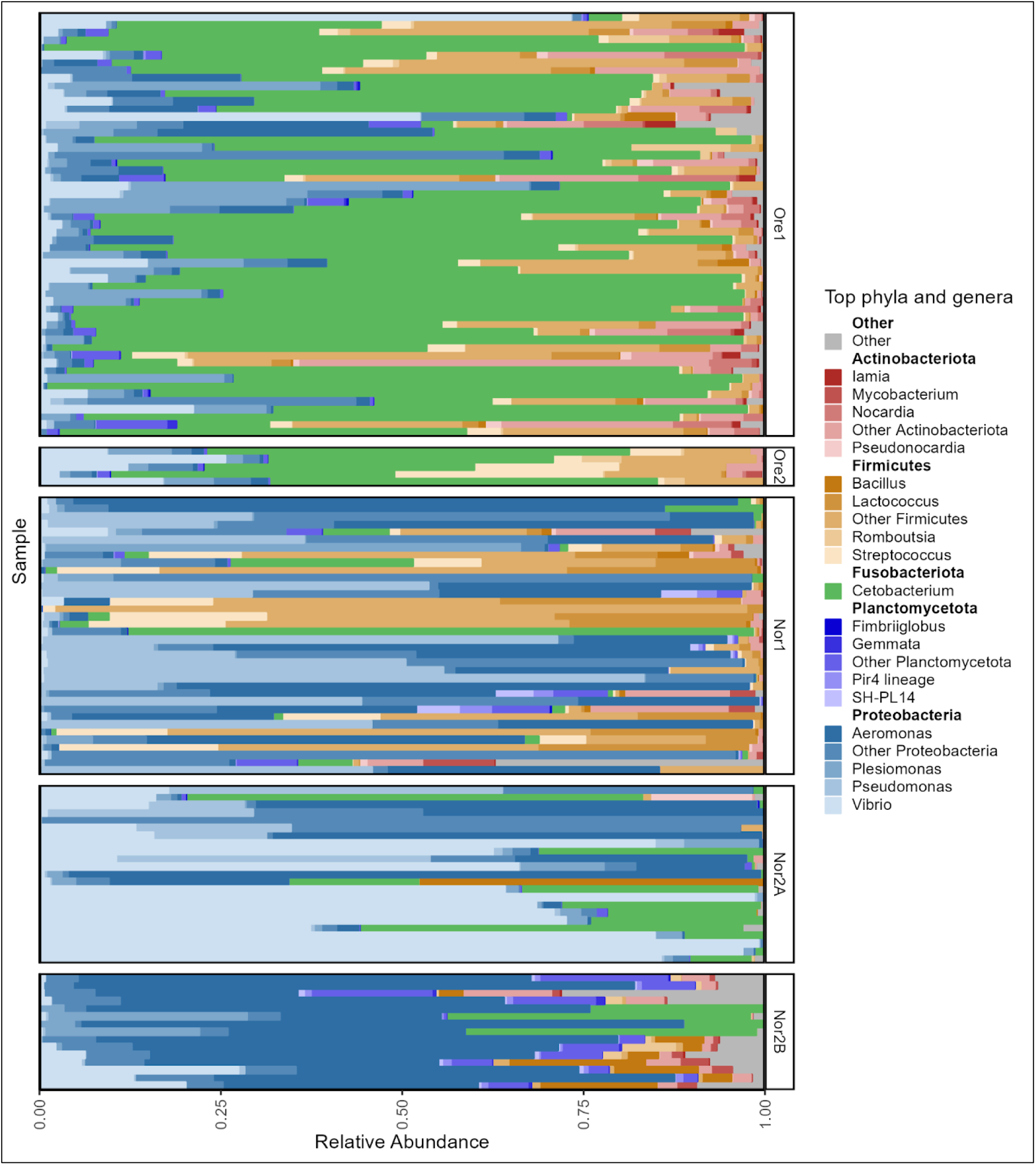
Phylum and genus-level taxonomic composition of zebrafish gut microbiome samples across aquaculture facilities. The mean relative abundance is grouped by aquaculture facility and is displayed for only the top five most abundant phyla and top four genera within each phylum as averaged across samples.

#### Aquaculture facility strongly influences zebrafish gut microbiome variation

Compositional differences in zebrafish microbiomes were visualized using a PCoA based on Bray-Curtis and unweighted UniFrac dissimilarity (Figure 7). Despite apparent overlap in the plot, the beta-diversity of zebrafish gut microbiomes varied significantly by both location and facility (PERMANOVA; Bray–Curtis and unweighted UniFrac: *p* < 0.001). Location explained 21.9% of variation in the Bray–Curtis and 12.3% in the unweighted UniFrac dissimilarities, while facility identity explained 11.5% and 10.2%, respectively. Genotypic status (WT vs. GM) appeared significant in the nested model (Bray–Curtis: *p* < 0.030; unweighted UniFrac: *p* < 0.001), but this effect did not remain when permutations were constrained within facility (Bray– Curtis: *p* = 0.167; unweighted UniFrac: *p* = 0.742), suggesting that, as in water, the apparent influence of genotype may be partly confounded with location or facility effects (Supplementary Figure 3). Pairwise PERMANOVAs confirmed significant variation in the composition of gut microbiomes between most facilities (*p*-adj. = 0.01), except for Ore1 and Ore2 (Bray-Curtis: *p*-adj. = 0.07, unweighted UniFrac: *p*-adj. = 0.01).

**Figure 7:**
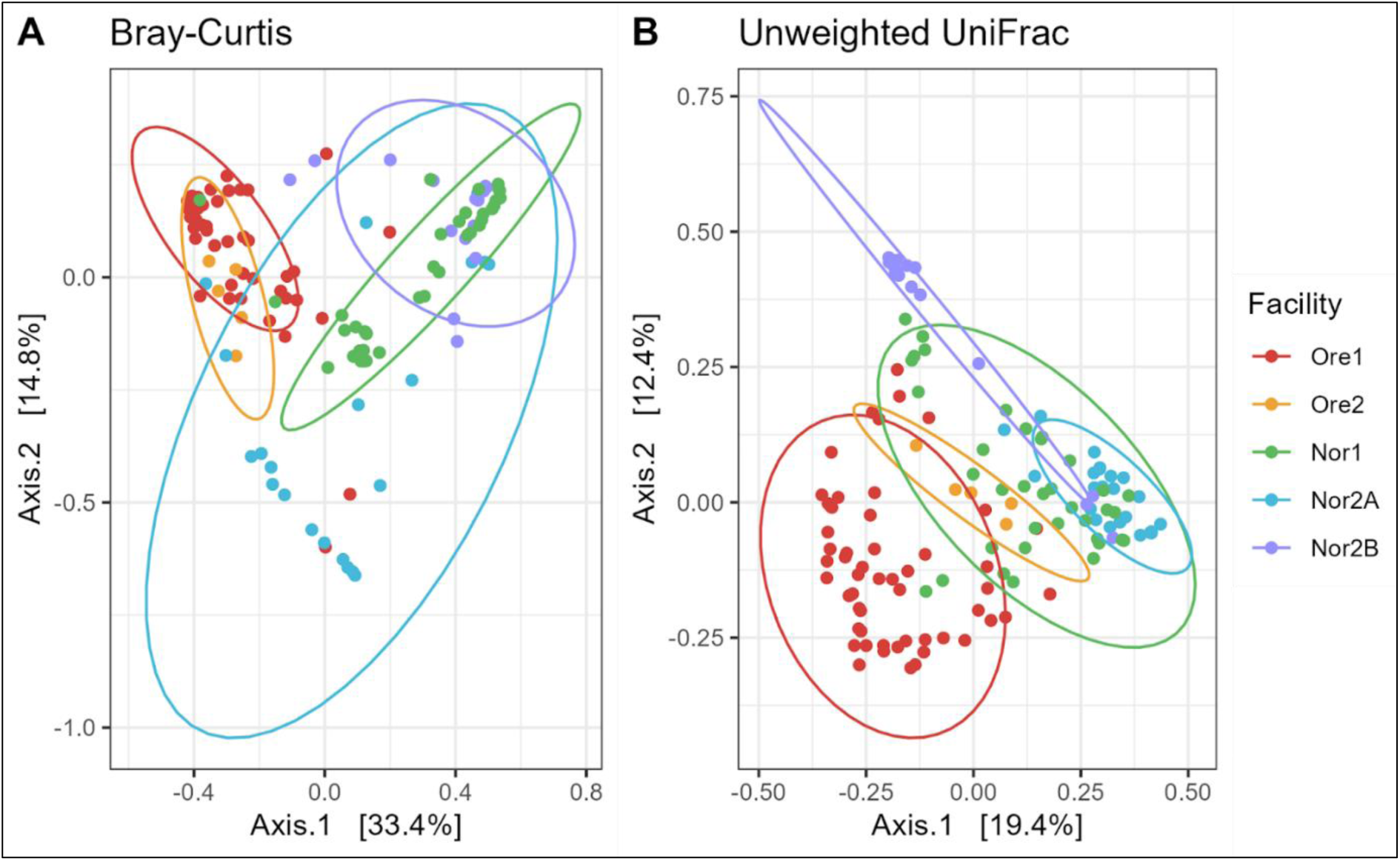
Beta diversity of zebrafish gut samples across aquaculture facilities. A principal coordinate analysis (PCoA) of (A) Bray-Curtis and (B) unweighted UniFrac dissimilarity. Colors represent facilities with normal confidence ellipses (95%).

The SIMPER analysis of fish microbiota revealed that facility differences were primarily driven by shifts in members of the Fusobacteria and Proteobacteria phyla, similar to the water microbiome patterns. *Cetobacterium* (Fusobacteria) was the dominant contributor to dissimilarity, accounting for 22-26% of the observed differences between Oregon and Norwegian facilities. *Vibrio* (Proteobacteria) followed as a secondary driver, contributing 20-23% of the dissimilarity in comparisons involving Nor2A, while *Aeromonas* (Proteobacteria) was particularly important in differentiating Nor2B from other facilities (27% contribution).

Distinctions among Norwegian facilities were also primarily driven by differential abundances of Proteobacteria taxa, with *Vibrio* and *Aeromonas* each contributing greater than 20% to the observed dissimilarities. These findings demonstrate that while similar bacterial phyla drive microbiome differences in both water and fish samples, the fish-associated microbiome shows more pronounced facility-specific signatures, particularly for Proteobacteria taxa that may reflect local environmental conditions or facility management practices (Supplementary Figure 4).

### Covariation Between Zebrafish Gut and Water Microbiomes

Once we confirmed significant variation in both fish and water microbiomes across facilities, we asked if there was significant covariance between fish and water microbiomes. We accomplished this by examining matched fish and water samples, meaning fish gut samples and water samples collected from the same tank, and measured their overlap in microbiome composition, defined as the extent of shared microbial taxa between the two sample types. We then used source tracking methods to estimate the proportion of water-associated taxa present in zebrafish guts and, conversely, gut-associated taxa present in tank water.

#### Fish and water microbiomes show greater similarity within than between facilities

Across all samples, 179 ASVs were shared between fish and water microbiomes, representing a subset of the 2,381 unique ASVs identified overall. Of these shared taxa, nearly half (81 ASVs) belonged to the Proteobacteria phylum, followed by Actinobacteriota (27 ASVs), Firmicutes (17 ASVs), Bacteroidota (14 ASVs), and Planctomycetota (12 ASVs). Many of these shared taxa were abundant in either fish or water (average relative abundance >1%), but just 12 were abundant in both. Of these, members of the *Cetobacterium*, *Pseudomonas*, and *Aeromonas* genera were shared at >5% average relative abundance in both fish and water.

We conducted pairwise comparisons of Bray-Curtis dissimilarity for fish and water collected from their own tank, tank water from elsewhere in the same facility, and water from tanks in other facilities (Figure 8). Overall, fish gut microbiomes were significantly more similar to both tank water from the same tank and within the same facility than tank water from other facilities (Kruskal-Wallis test: *p* < 0.001; pairwise Wilcoxon test: same tank vs. same facility, *p*-adj. = 0.013; same tank vs. other facility, *p*-adj. < 0.001; same facility vs. other facility, *p*-adj. < 0.001). Consistently, samples of paired fish and water microbiomes from either the same tank or the same facility were significantly more similar than fish and other facility water microbiomes. However, this trend was not observed in Nor1 (Supplementary Figure 5). The similarity between fish and water samples in the Oregon facilities was significantly greater than that seen in the Norwegian facilities.

**Figure 8:**
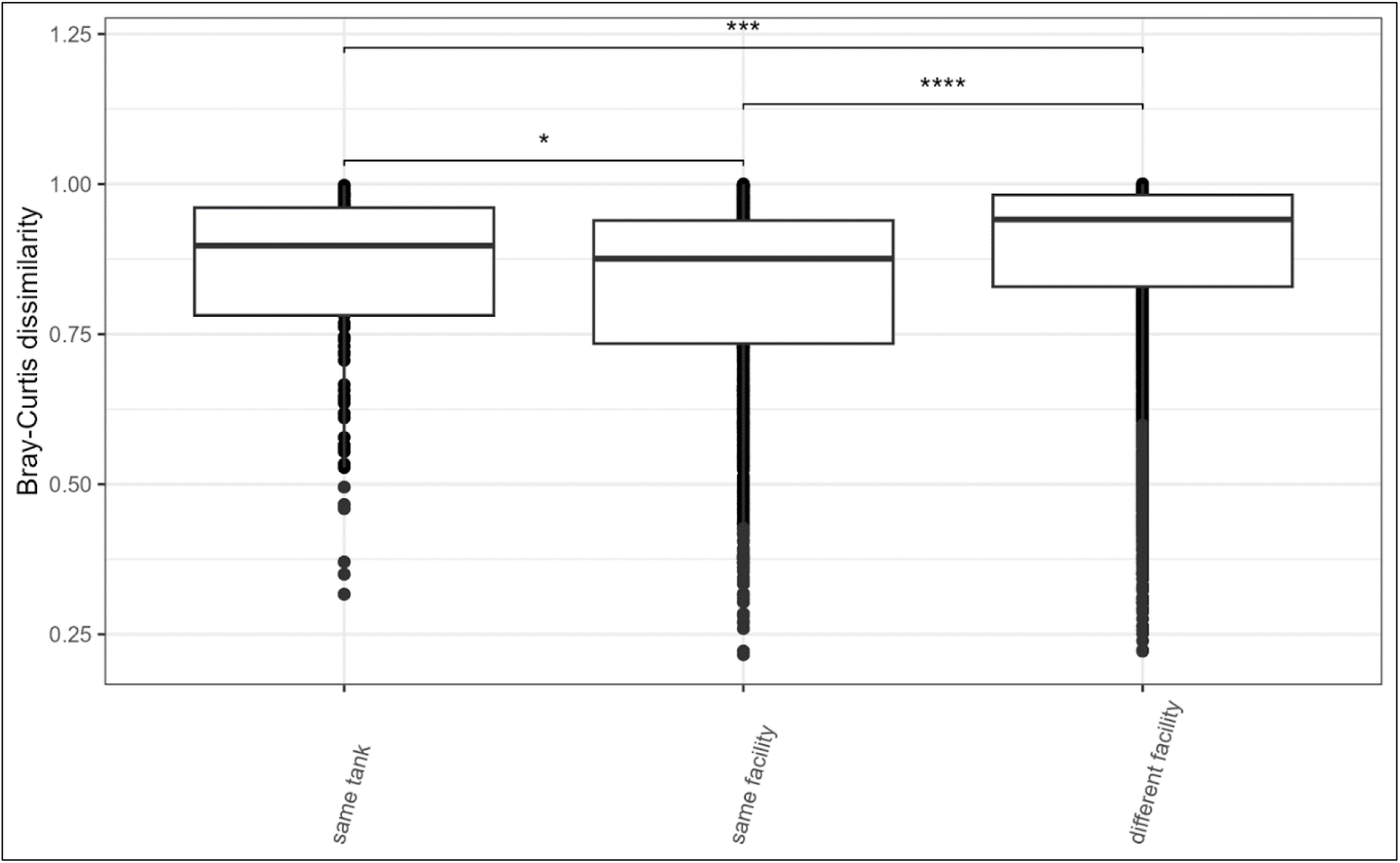
Tukey-style box and whisker plot featuring pairwise beta-diversity values using Bray-Curtis distance for fish gut microbiome samples paired with either 1) water from the same tank (‘same tank’), 2) water from tanks within the same facility but not in the same tank (‘same facility’), or 3) water from tanks in other facilities (‘different facility’). Significance calculated with Kruskal-Wallis and pairwise Wilcoxon rank-sum test with Benjamini-Hochberg p-value correction (* = p < 0.05, ** = p < 0.01, *** = p < 0.001, **** = p < 0.0001, ns = non-significant).

#### The influence of the water microbiome on zebrafish gut microbiomes varies by facility

To assess the extent to which the tank water microbiome contributes to the zebrafish gut microbiome, we used Fast Expectation-Maximization Microbial Source Tracking (FEAST), a probabilistic modeling approach that estimates the proportion of microbial taxa in a given sample (the “sink”) that can be attributed to potential source environments [20]. This method is particularly useful for identifying directional influences in microbial transmission, helping to determine the extent of environmental shaping on host-associated microbiomes.

In this analysis, zebrafish gut microbiomes were first classified as “sinks” and water microbiomes as “sources” to estimate the proportion of gut bacteria potentially acquired from the surrounding tank environment. Because microbial exchange can be bidirectional, we also performed the reverse analysis, treating fish as the source and water as the sink, to assess whether microbial taxa from zebrafish were being introduced into the water column.

The results indicated significant variation in the proportion of water-derived microbiota in zebrafish guts across facilities (Kruskal-Wallis: *p* < 0.001). Based on relative abundance estimates, fish from the Nor2A and Ore2 facilities exhibited significantly higher proportions of tank water microbiota in their guts compared to those from Ore1, Nor1, and Nor2B (Figure 9A).

**Figure 9:**
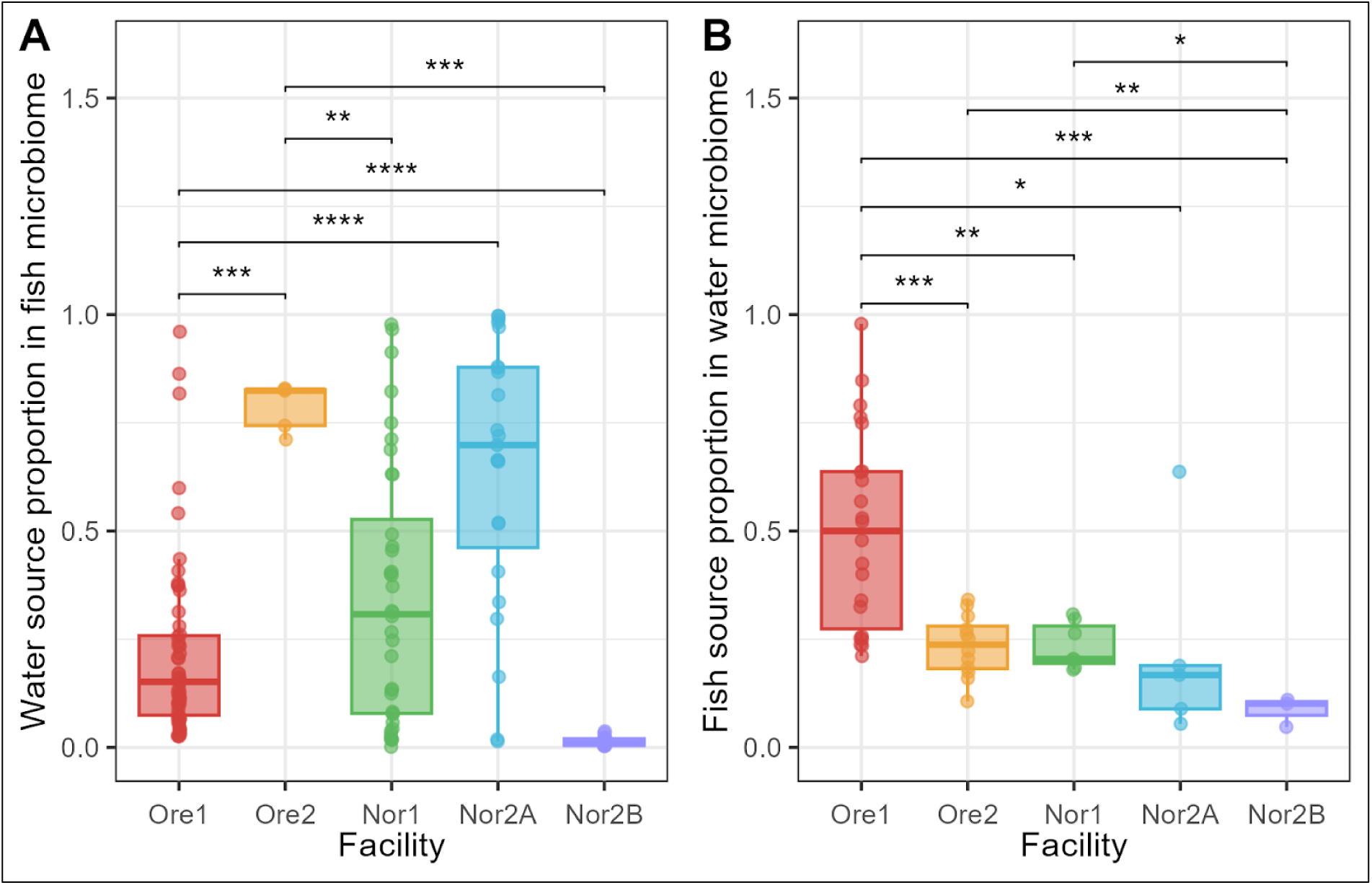
Source proportions for (A) water microbiota found in the zebrafish gut microbiome and (B) fish microbiota found in water microbiomes. Proportions were calculated using probabilistic models within the R package ‘FEAST’. Significance calculated with pairwise Wilcoxon rank-sum test with Benjamini-Hochberg p-value correction (* = p < 0.05, ** = p < 0.01, *** = p < 0.001, **** = p < 0.0001, ns = non-significant).

Conversely, in the reverse analysis, the Ore1 and Nor2B facilities were the only sites where fish microbiota were found in greater proportions in water than water microbiota in fish (Figure 9B). Notably, Nor2B fish consistently contained the lowest proportion of shared microbiota, indicating that the strength of environmental influence on gut microbial composition varies by facility.

## Discussion

In this study, we aimed to characterize microbial variation across aquaculture facilities and assess the extent of microbial sharing between zebrafish gut and water microbiomes. By analyzing paired tank water and zebrafish gut microbiomes from five research-focused aquaculture facilities, we found that facility-specific factors strongly shape microbial community composition in both fish and water. While water and fish microbiomes remain compositionally distinct, the most abundant microbial taxa are widely shared between the two. Notably, zebrafish gut microbiomes exhibit greater similarity to water microbiomes within their own facility than to those from other facilities. Source tracking analyses further suggest that the water microbiome influences the zebrafish gut microbiome, but the strength of this interaction varies across facilities. Reciprocal analyses indicate that, in some cases, microbial exchange may be bidirectional, with fish also contributing to the surrounding water microbiome.

### Variation in Facility Water Microbiota

Beta-diversity analyses of the facility water microbiomes revealed clear differences in microbiome composition between locations (Oregon vs. Norway) as well as in the variation among facilities within a geographic location. Examining the community composition provides some clues to the driving forces behind these differences. Across all facilities, two phyla— Proteobacteria and Fusobacteriota—accounted for more than 75% of total reads in most samples. While Proteobacteria were represented by multiple genera, Fusobacteriota was almost exclusively represented by a single genus, *Cetobacterium*, assigned to a single ASV (ASV1). The Oregon facilities, in particular, were dominated and differentiated by the relative abundance of *Cetobacterium*.

*Cetobacterium* has been identified as a “core” member of the zebrafish gut microbiome [14, 15] as well as in a wide range of other freshwater fish species [21–23]. Consistent with this host-association, our data showed *Cetobacterium* at substantially higher relative abundances in fish samples than in water samples across all facilities. In zebrafish, *Cetobacterium* appears to be enriched in the intestines of adults [24] and has been associated with potential host benefits, including glucose homeostasis, reduced parasite burden, and probiotic effects on liver health [25–27]. This evidence strongly suggests that the *Cetobacteria* found in the water samples are likely host-associated microbes that originate from, or are at least strongly amplified by, the zebrafish hosts in this study system.

Other highly prevalent taxa in the Oregon facility water samples, including *Psychrobacter* and *Vibrio*, are also widely associated with fish microbiomes [14, 28]. While some *Vibrio* species are known pathogens [30], this diverse genus includes many species that are common associates of healthy fish and aquatic environments, with some even showing potential probiotic properties in fish [31]. This pattern is similar to how *E. coli* can be both commensal and pathogenic in humans-the genus encompasses both beneficial and harmful members. Given that source tracking analyses indicated that fish microbiota contributed up to 50% of the water microbiome in Ore1 and 25% in Ore2, it is likely that microbial inputs from zebrafish play a significant role in shaping the water microbiome in these facilities.

In contrast, the Norwegian facilities were dominated by Proteobacteria, though the most abundant taxa varied by facility. Many of these taxa are commonly found in open water and sediments, yet their association with fish appears to be primarily in the context of stress, disease, or experimental manipulation rather than as normal members of healthy fish microbiomes [32]. For instance, in Nor2B, the only flow-through system represented in this study, dominant taxa included *Acidovorax* and *Nevskia*, which are primarily associated with wastewater and open water environments [33, 34]. While *Acidovorax* has been identified as a potential contaminant in some microbiome studies [35], it is also documented as a fish-associated taxon and was not flagged as a contaminant in our decontam analysis. Notably, *Nevskia* has been enriched in larval zebrafish exposed to microplastics [36], suggesting that environmental stressors may influence its abundance.

Nor1 also had abundant taxa such as *Delftia*, which is typically found in water and soil but has occasionally been observed in pathogen-challenged zebrafish guts [37–40]. Meanwhile, Nor2A was dominated by *Rheinheimera*, a taxon frequently detected in aquatic systems and soil [41, 42] and enriched in zebrafish exposed to antibiotics, particularly in germ-free derivation processes [43, 44]. This pattern suggests that the Norwegian facilities may harbor microbiomes more reflective of environmental colonization or stress responses rather than potentially more host-adapted communities observed in the Oregon facilities.

One possible explanation for the striking difference between the Oregon and Norwegian facility water microbiomes is the relative age of the facilities and the total number of housed fish. The Oregon facilities have been in operation longer than those in Norway-Ore1 and Ore2 were established in 2012 and 2001, respectively, whereas Nor1 opened in 2014 and Nor2A/Nor2B in 2015. Since these facilities use recirculating aquaculture systems (RAS), the water in the Oregon facilities has been recirculating for significantly longer. Additionally, the Oregon facilities house far more fish (approximately 50,000 in Ore1 and 30,000 in Ore2) compared to those in Norway (approximately 10,000 in Nor1 and 100–200 in Nor2A and Nor2B).

While biofilters would be considered ‘mature’ after nearly a decade of operation, the longer operational period combined with higher fish populations in the Oregon facilities may have allowed for greater cumulative microbial inputs from operational factors such as feeding regimes, cleaning routines, human activity, and fish waste product. Over time, these accumulated inputs may shift the microbial communities within the biofilter, water column, and biofilms throughout the facility plumbing to become increasingly reflective of the fish microbiome rather that the source water or initial environmental colonizers. Similar patterns have been observed in built environments, such as offices and homes, where the prolonged presence and activity of hosts shape the environmental microbiome through cumulative biological inputs [45, 46].

Additionally, the Norwegian facilities house more disease-model zebrafish genotypes compared to the Oregon facilities. Disease-model and immunocompromised zebrafish may have reduced host selection capacity, making them more susceptible to colonization by environmental microbes that would typically be excluded by host immune filtering. For example, genetically modified lines with mutations affecting ciliary function, such as the *foxj1b* mutant lines housed at Nor1 (Supplementary Table 6), can compromise host defenses. In zebrafish, the *foxj1b* gene is essential for motile cilia formation [47], and its mammalian homolog *Foxj1* also regulates adaptive immune responses by repressing inflammatory pathways [48]. Compromised ciliary and potentially immune function may reduce the host’s ability to selectively maintain host-associated microbiota, allowing environmental microbes to colonize more readily. Consequently, the greater prevalence of pathogen- and antibiotic-associated microbiota in Norwegian facility water could be influenced not only by the higher proportion of disease-model zebrafish housed in these facilities, but also by the compromised ability of these fish to selectively maintain host-adapted microbial communities.

One hypothesis about what drives variation in microbiome composition among facilities is variation in bacterial growth rates due to differences in water exchange rates of the tanks. It has been proposed that shorter hydraulic retention times in RAS systems, which are directly tied to higher water exchange rates in tanks, promote r-selected bacterial growth whereas longer retention times promote K-selected bacterial growth [49]. Although we did not directly measure water exchange rates, most facilities used similar zebrafish husbandry equipment—Ore1, Ore2, Nor1, and Nor2A all utilized Techniplast (Buguggiate, Italy) zebrafish rearing systems. Given that variation in water exchange rates in these systems is constrained by tank design, variation among tanks within a facility is likely greater than the variation in average tank water exchange rates between facilities. Therefore, water exchange rate alone is unlikely to drive the observed differences in microbial composition among facilities.

Another more plausible driver of the observed facility-level differences is variation in tank maintenance and cleaning protocols. The Oregon facilities maintained the strictest cleaning protocols, changing tanks every two weeks or earlier if visible algal or biofilm growth was observed (T. Mason, Ore1 facility manager, personal communication, January 30, 2021; D. Lains, Ore2 facility manager, personal communication, September 21, 2022). The Norwegian Nor2A and Nor2B facilities followed a similar two-week cleaning schedule (F. Jutfelt, Nor2A/Nor2B facility manager, personal communication, January 12, 2024), whereas Nor1 changed tanks much less frequently—approximately once every two to three months (E. Yaksi, Nor1 facility manager, personal communication, June 14, 2022). All facilities used personal protective equipment to limit potential cross-contamination.

Despite the differences in cleaning protocols between Nor1 and Nor2A/B, nearly all sampled tanks in the Norwegian facilities exhibited at least some visible biofilm or algal growth, while the Oregon facility tanks appeared largely free of biofilms. This observation suggests that cleaning intensity, rather than frequency alone, may be a key factor driving differences in microbial composition between facilities.

The more intensive cleaning regimen in the Oregon facilities likely created distinct ecological conditions that favored fish-associated microbiota in the water column. Frequent disturbance through aggressive cleaning would reduce overall microbial biomass and remove established, competitive bacterial communities (i.e., K-selected taxa) in the tank water. This disturbance could promote the detection of fish-associated microbes through two potential mechanisms: first, fish-associated microbes dispersing from hosts could rapidly colonize the newly available niche space in the water column; alternatively, these microbes could be detected simply due to the reduced background water microbial biomass, making fish-derived taxa more prominent in the relative abundance profiles without necessarily requiring growth in the water. Either mechanism could explain the strong influence of fish microbiota observed in the Oregon water samples.

In contrast, the greater biofilm and algal growth observed in Norwegian facility tanks may provide competitive taxa that limit the establishment of fish-associated microbes in the water column. If these biofilms help stabilize the water microbiome, this could explain why Norwegian facility water samples were more compositionally similar to one another than those from Oregon.

Ultimately, the observed differences in water microbiomes across facilities are likely shaped by multiple interacting factors, including, but not limited to, facility age, fish density, microbiota dispersal from zebrafish, tank maintenance routines, and microbial competition within the water column. Given these differences, it follows that zebrafish reared in different facilities experience different microbial environments, which could, in turn, shape the composition of their microbiomes. The next section explores how these facility-level differences translate to variation in the zebrafish gut microbiome.

### Variation in the Zebrafish Gut Microbiome

The zebrafish gut samples shared many similarities to patterns seen in the tank water samples. Although variation in alpha-diversity between facilities was far greater, the composition of the zebrafish microbiome showed significant influence of both location and facility origin. In Roeselers et al. (2011), the authors compared the microbiomes of wild and domesticated zebrafish. In their paper, Proteobacteria represented over 99% of all reads in wild-caught zebrafish raised in laboratory aquaculture facilities but made up considerably smaller proportions of the microbiome of zebrafish genotypes that had been maintained for many generations in such facilities. Notably, this pattern was seen in both the Nor2A facility and the Nor2B facility, where Proteobacteria comprised the greatest proportion of reads-upwards of 95% and 75%, respectively-while making up a far lesser proportion of reads in the other facilities. Unlike the other facilities, which maintain highly domesticated zebrafish lines like AB (likely 50+ generations removed from wild), the Nor2B zebrafish are approximately sixth generation descendants of wild zebrafish collected in India in 2015. Those in Nor2A are a cross between these wild descendants and a domesticated Casper line. This proximity to “wildness” could explain some of the compositional patterns seen in these facilities. However, both the Fusobacteriota that dominates the Oregon facilities and the Proteobacteria abundant in the Norwegian facilities are considered part of the shared ‘core’ microbiome [14, 15]. Effects of genotype on the composition and structure of the host microbiome have been well documented in zebrafish research [50]. As such, we attempted to include genotype as a factor in our mixed models. In the facilities, there were many different lines and sublines of zebrafish that could be considered either wild-type or a genetic modification of these lines and were often confounded with facility (i.e. wild-type strain ‘ABC’ was only found in Ore1 while the genetically modified strain ‘Elipsa’ was only found in Nor1). Therefore, we created a binary classification where all lines that could be considered wild type were coded as ‘WT’ and all lines that were genetically modified for a specific experimental purpose as ‘GM’ (Supplementary Figure 6). This factor was not significant in our constrained model. However, genotype complexity—introduced by decades of cross-breeding and genetic modifications [51, 52]—may still play a role in structuring the microbiome, warranting further investigation.

### Zebrafish Gut and Water Covariation

Lastly, to explore the variation observed in the microbial communities of both fish and water, we found that many of the patterns in diversity and composition unique to individual facilities were reflected in both sample types. This suggests active sharing of microbiota between the two. However, it is difficult to establish whether this sharing is reciprocal or primarily unidirectional. To investigate the directionality of transfer, we employed two complementary approaches. First, we considered the identity of the shared bacterial taxa and whether they were considered predominantly aquatic or fish-associated in the literature. A key caveat to this approach is that taxonomic associations can be highly context-dependent, varying by environment, host species, or experimental conditions. Our compositional sequencing approach cannot directly assess microbial function or the nature of host-microbe interactions, so inferences about directionality based on reported associations should be interpreted cautiously [53, 54].

To complement this approach, we used a source tracking method to estimate whether the shared taxa were more likely to have originated from the water or the fish gut. The source tracking analysis, conducted with the R package ‘FEAST’, assumes directionality in the transmission of microbiota. Given the lack of prior conclusive evidence for the direction of transmission between fish and water in these systems, we performed the source tracking analysis twice: once with fish designated as the ‘source’ and once with fish as the ‘sink’. This reciprocal analysis aimed to identify any clear patterns of directionality, such as a large proportion of water microbiota found in fish, but minimal fish microbiota found in water.

In the Oregon facilities, we observed a significant fraction of ASVs shared between fish and water samples, with the water community dominated by taxa from the phyla Fusobacteriota and Firmicutes. As previously discussed, the only genus identified in Fusobacteriota- *Cetobacterium*- is widely associated with freshwater fish [14, 22]. Many abundant Firmicutes taxa showed similar fish associations. For instance, *Epulopiscium* and *Exiguobacterium* have documented in fish gut and skin samples, respectively [55, 56]. This prevalence of fish-associated microbiota in the water microbiome suggests substantial microbial input from the fish hosts, or alternatively, reflects the reduced overall biomass in the water column that increases the relative detection of fish-derived taxa (as discussed earlier).

The Norwegian facilities presented a contrasting pattern, with fewer abundant water taxa also found at high abundances in the fish gut microbiome. This divergence likely reflects the more stable water microbiome resulting from reduce cleaning-induced disturbance, while fish hosts maintain selective pressure on their microbial communities-though this selectivity can vary with genotype [57], immune/health status [58], and environmental stressors [59]. Notably, dominant members of the water microbiome were not necessarily found at high abundances in the fish microbiome. While relative Proteobacteria proportions appeared similar between water and fish across all facilities, highly abundant water microbiota did not correlate with proportionally greater abundance in the zebrafish microbiome. Instead, facility-level variation in the fish microbiomes was primarily driven by compositional shifts among microbial taxa already considered part of the normal zebrafish gut microbiome. However, zebrafish and water did share rare ASVs unique to each facility, indicating that beyond shifts in numerically dominant taxa, facility-specific rare taxa may also contribute to the observed differences. This pattern mirrors observations in fish microbiomes across environmental gradients, where rare taxa distinguished microbial communities between conditions despite low numerical abundance, suggesting these taxa may play disproportionately important ecological roles [60].

Source tracking results further highlighted the highly variable nature of microbial sharing between fish and water across different aquaculture facilities. (Members of the water microbiome were found to be more prevalent in the gut microbiota of fish reared in RAS systems (Ore1, Ore2, Nor1, and Nor2A) compared to the flow-through system (Nor2B). However, since we only investigated one flow-through facility, these results may not be representative of all facilities using flow-through systems. Additionally, microbial source tracking revealed no inverse relationship between the proportion of water-derived microbiota in fish and fish-derived microbiota in water across facilities. Facilities showing smaller contributions of water microbiota to fish microbiomes did not correspondingly exhibit larger contributions of fish microbiota to water microbiomes. For example, facility Nor2B showed relatively low proportions of both water-to-fish and fish-to-water microbial contributions ∼5% and 10%, respectively, suggesting substantial microbial inputs from untracked sources. In contrast, the Ore1 facility exhibited the largest proportion of fish-derived microbiota in its water samples (∼50%), far exceeding all other facilities. Excluding Ore1, fish microbiota contributions to water communities remained relatively consistent across facilities, suggesting fish provide a baseline microbial input to their environment. However, the influence of water microbiomes on fish communities varied substantially by facility, likely driven by the same operational and system-specific factors that contributed to water microbiome variation across facilities.

These findings align with our observations of the relationship between the zebrafish and water microbiomes across facilities. Variations in the water microbiome do not appear to drive corresponding compositional variations in the zebrafish microbiome, but there is clear evidence of microbial sharing between the two. This is consistent with observations in other aquatic organisms, such as common bottle-nose dolphins, California sea lions, Atlantic cod, and Mediterranean gilthead sea bream. In these cases, while host and environmental microbiomes share some similarities, the abundance and diversity of environmental microbiota do not necessarily correlate with variation in the host microbiome, and the microbiomes of each remain largely distinct [28, 61, 62].

A significant limitation to this study is that the microbiomes of fish and water were sampled at a singular time point. Previous work [63] shows that the water microbiome within facilities varies significantly over time, and since this variation is correlated with changes in the fish gut microbiome, this temporal variation may influence the relative influence of the water microbiome on the zebrafish microbiome. Future research that includes repeated sampling over time could provide more accurate insight into these interactions.

Although we cannot fully define the mechanisms or directionality of transmission between the zebrafish gut microbiome and surrounding water microbiome, our results demonstrate that zebrafish and water microbiomes share a large proportion of microbiota. Both microbiomes are heavily influenced by aquaculture facility and geographic location. Zebrafish are widely used as model organisms in basic research, and variation in their microbiome can significantly impact the reproducibility of experimental outcomes. As noted earlier, the host-microbiome relationship mediates many aspects of host health and function, and previous studies in mice and primates have linked microbial variation to disparate experimental results associated with facility origin [7, 10, 64]. The microbiome variation we observed across facilities and regions could potentially reduce the reproducibility of zebrafish research focused on microbiome-mediated processes. One possible driver of this variation is differences in hygiene practices across facilities. Other factors, such as variations in fish diet, water quality, and stocking density, could also contribute to differences in microbial communities. Improving consistency in hygiene practices, along with controlling other facility-specific factors like diet and environmental conditions, could enhance reproducibility. Future work should aim to identify and standardize such factors across zebrafish facilities, ultimately reducing microbiome variation and improving experimental consistency.

## Conclusions

Our results demonstrate that even in relatively controlled aquaculture facilities, facility-specific differences can significantly impact both water and zebrafish gut microbial communities. We found evidence that variation in the zebrafish microbiome is driven, in part, by the environmental acquisition of microbiota from the tank water, although the magnitude of this influence varies across facilities. Microbial sharing between zebrafish and their environment likely represents just one aspect of a broader array of facility-specific factors that shape the zebrafish gut microbiome. Future studies should aim to isolate and characterize these factors to better understand the mechanisms and extent of microbiome transmission. This study expands our current understanding of the factors influencing the microbiome of zebrafish, a widely used animal model, and underscores the need for more in-depth sampling and monitoring to identify the key drivers of microbiome assembly in aquaculture facilities. Furthermore, we caution that when using zebrafish models sourced from single facilities, researchers should be mindful of facility-specific microbiome variation, as it could greatly influence experimental outcomes.

## Methods

### Study Sites

This study was conducted at five zebrafish aquaculture facilities spanning two geographic regions: three located on the University of Oregon campus in Eugene, Oregon, USA, and two located at the Norwegian University of Science and Technology (NTNU) in Trondheim, Norway. These facilities were chosen because they provide a unique system to investigate host– environment microbial interactions across spatial scales and facility designs. Specifically, their close clustering within each geographic region allows investigation of variation in microbiome composition both within and between facilities in close proximity, while the inclusion of facilities across continents introduces larger-scale geographic comparison. Additionally, the sites encompass a range of rearing approaches, genotypic diversity, and life support systems representative of common practices in modern aquaculture, making the results broadly applicable to other research settings.

Both U.S. facilities included in this study are housed on the main University of Oregon campus. The first, the Aquatic Animal Care Services Zebrafish Facility (Ore1), was established in 2012 and originally stocked with fish from commercial suppliers. It spans approximately 1,000 square meters with an average capacity of 50,000 fish (maximum capacity ∼88,000) representing around 2,900 genotypic strains. Fish are housed in 3.5-liter acrylic tanks connected to two closed-loop recirculating aquaculture systems (RAS) that utilize automated mechanical and biological filtration and ultraviolet (UV) disinfection. There is also a separate quarantine area with a dedicated flow-through system.

The second U.S. facility, the Zebrafish International Resource Center (Ore2), was built in 1999 and began operation in 2001. Ore2 covers 900 square meters and maintains an average of 30,000 fish (maximum capacity ∼150,000), representing approximately 45 live genotypic lines and over 44,000 genotypes stored in cryogenic preservation. The layout and life support systems in Ore2 are comparable to those in Ore1, with fish housed in both 3.5-liter acrylic and 75-liter glass tanks. This facility is located approximately 300 meters from Ore1, enabling comparisons between adjacent facilities that share a similar local environment but operate independently. The three Norwegian zebrafish facilities are located on the NTNU Øya and Gløshaugen campuses in Trondheim, Norway. The Yaksi Lab facility (Nor1) is situated on the Øya campus and was built in 2014 for research on brain function and neurological disease. The facility is 70 square meters in size and houses approximately 10,000 zebrafish representing over 100 genotypic strains. Fish are kept in 3.5-liter acrylic tanks attached to a RAS with UV-treatment. There is also a separate quarantine area using a flow-through system with roughly 50 tanks.

The Jutfelt Lab facility (Nor2) is located on the Gløshaugen campus and was originally established as a general animal facility in the 1990s before being repurposed to house fish between 2015 and 2020. The lab maintains both zebrafish and guppies (*Poecilia reticulata*) in five separate rooms (each approximately 10–30 square meters) with independent life support systems. While administratively considered one facility, we reference two zebrafish-only rooms within the Jutfelt Lab as Nor2A and Nor2B due to their distinct system configurations.

Nor2A contains approximately 50, 3.5-liter acrylic tanks on a standalone RAS with UV-treatment. Nor2B houses 14, 75-liter glass tanks connected to a flow-through filtration system with no UV-treatment. Together, the Nor2 rooms house roughly 2,000 fish. Unlike the other facilities, which maintain common laboratory zebrafish lines, the Jutfelt Lab exclusively houses zebrafish descended from wild-caught individuals collected in India in 2016 (∼6–10 generations removed).

### Sample Collection and Processing

#### Tank water

We collected tank water from a representative number of tanks within each facility, targeting tanks connected to a single RAS system and housing adult zebrafish. We adjusted the number of tanks sampled based on the size and housing configuration of each facility (n = Ore1 (27); Ore2 (17); Nor1 (7); Nor2A (6); Nor2B (3)). For each RAS system, we chose one to two adjacent shelf sets, then randomly selected three shelves from each set. On each selected shelf, we randomly sampled three tanks. When fewer than three shelves or tanks were available (e.g., Nor2B, which had two shelves with two tanks each), we sampled all available tanks. To collect microbial biomass, we passed 150 mL of tank water through a Sterivex filter cartridge (MilliporeSigma, Massachusetts, USA) using a 60-mL syringe. We capped the Sterivex filters and kept them on ice until processing. If we could not perform DNA extraction immediately, we stored the dry filter cartridges at −80 °C.

#### Zebrafish

We collected zebrafish from the same tanks used for water sampling (n = Ore1 (55); Ore2 (5); Nor1 (36); Nor2A (23); Nor2B (15)). We euthanized fish via rapid chilling and sampled four individuals per tank. If fewer than four fish were available or sampling was limited, we collected as many individuals as possible.

On the same day, we aseptically dissected the whole intestine from each fish and transferred it into a 2-mL RHINO screw-cap tube containing zirconium oxide beads and 400 µL enzymatic lysis buffer (20 mM Tris-Cl, pH 8.0; 2 mM sodium EDTA; 1.2% Triton X-100). If immediate DNA extraction was not possible, we stored the tubes at −80 °C.

### Microbial DNA Extraction

#### Tank water

We performed DNA extraction using the DNeasy PowerWater Sterivex kit (QIAGEN, Carlsbad, California, USA). DNA was extracted following the standard protocol provided in the kit handbook (2009, pg. 9).

#### Zebrafish

We performed DNA extraction from zebrafish gut samples using an adapted protocol based on the DNeasy Blood & Tissue Kit Quick-Start protocol, as described in Stephens et al. (2015).

### 16S rRNA Gene Amplification, Library Preparation, and Sequencing

DNA extraction products were quantified using a Qubit dsDNA HS Assay Kit (Thermo Fisher Scientific, Waltham, MA, USA). To amplify the V4 region of the 16S rRNA gene, dual-indexed primers (515F–806R) were used, with unique 8-bp index sequences incorporated as described in Caporaso et al. (2011): 515F (AATGATACGGCGACCACCGAGATCTACAC[index]TATGGTAATTGTGTGCCAGCMGC CGCGGTAA) and 806R (CAAGCAGAAGACGGCATACGAGAT[index]AGTCAGTCAGCCGGACTACHVGGGTWT CTAAT).

PCR reactions included 12.5 µL NEBNext Q5 Hot Start HiFi PCR Master Mix (New England Biolabs, Ipswich, MA, USA), 10.5 µL DNA template, 1 µL Bovine Serum Albumin (Thermo Fisher Scientific), and 1 µL of mixed primers (12.5 mM). Cycling conditions were: initial denaturation at 98 °C for 30 seconds; 30 cycles of 98 °C for 10 seconds, 61 °C for 20 seconds, and 72 °C for 20 seconds; and a final extension at 72 °C for 2 minutes. Samples were held at 4 °C until removal from the thermocycler.

To confirm amplification success, PCR products were visualized on an agarose gel. Amplicon libraries were cleaned twice using Mag-Bind RxnPure Plus isolation beads with a modified 0.8× reaction volume protocol (Omega Bio-Tek, Norcross, GA, USA). Concentrations were then quantified using the Quant-iT 1× dsDNA HS Assay Kit (Thermo Fisher Scientific) and a SpectraMax M5E Microplate Reader (Molecular Devices, San Jose, CA, USA). Libraries were pooled at equimolar concentrations and sequenced at the University of Oregon Genomics & Cell Characterization Core Facility on an Illumina MiSeq (Illumina, San Diego, CA, USA) with 150-bp paired-end reads.

### Bioinformatics

We performed all bioinformatics processing in ‘R’ [65]. We demultiplexed and then denoised the sequences to construct an amplicon sequence variant (ASV) table using DADA2 1.16 [66]. We assigned taxonomy to sequences using the RDP Classifier and Silva NR99 v.138.1 16S rRNA gene reference database [67, 68]. We evaluated sample contaminants using the prevalence method with ‘decontam’, which identifies contaminants based on their frequency of occurrence in negative control samples relative to true samples [69]. This analysis identified two ASVs as contaminants: ASV 226 (*Janthinobacterium*) and ASV 372 (unspecified Gemmataceae), which were subsequently removed from all downstream analyses.

### Statistics

We conducted all statistical analyses in ‘R’ (R Core Team, 2024). Alpha diversity metrics (Shannon and Inverse Simpson) and beta diversity metrics (Bray-Curtis and unweighted UniFrac) were calculated using the ‘vegan’ package and ASV tables [70, 71]. All plots were created with the ‘ggplot2’ package [72].

To test for differences in alpha diversity between water and zebrafish gut microbiomes across facilities, we applied non-parametric Kruskal-Wallis tests [73], followed by pairwise Wilcoxon rank-sum tests with Benjamini-Hochberg p-value correction [74, 75]. To assess the effects of geographic location, facility, and genotype status on microbial community dissimilarity, we performed hierarchical permutations using permutational multivariate analysis of variance (PERMANOVA) with the ‘adonis2’ function in the ‘vegan’ package. We used a nested design accounting for the hierarchical structure of the data, where facilities are nested within geographic locations (Norway vs. Oregon). Permutations were constrained to occur within the appropriate hierarchical levels to avoid pseudoreplication: facility-level effects were tested by permuting samples within locations, while location effects were tested by permuting whole facilities. Genotype status effects were tested using unrestricted permutations within each facility. All analyses used Bray-Curtis and unweighted UniFrac dissimilarity matrices and 9999 permutations. Pairwise PERMANOVA tests were performed using the ‘pairwise.Adonis’ package [76], and principal coordinate analysis (PCoA) was used to visualize microbiome composition clustering.

To generate relative abundance plots, ASVs were first agglomerated by taxonomic classification level. Relative abundances were then calculated by dividing the sum of ASV counts per taxon by the total number of counts per sample. These values were visualized using nested bar plots created with the ‘fantaxtic’ package [77].

To identify ASVs contributing most to differences between fish and water microbiota across facilities, we performed a Similarity Percentage (SIMPER) analysis using Bray-Curtis dissimilarity [19]. This method partitioned Bray-Curtis dissimilarity and calculated the average contribution of each taxon to compositional differences between sample pairs.

For pairwise beta-diversity comparisons, Bray-Curtis dissimilarity values were grouped into three spatial categories: (1) fish and water samples from the same tank, (2) from different tanks within the same facility, and (3) from different facilities. Dissimilarity values were plotted by group, and differences were tested using Kruskal-Wallis and pairwise Wilcoxon rank-sum tests with Benjamini-Hochberg correction.

To evaluate microbial exchange between fish guts and surrounding water, we applied Fast Expectation-Maximization for Microbial Source Tracking [20]. Within each facility, water samples were assigned as potential sources and fish samples as sinks. Source proportion values were summed across all input sources for each sink and visualized as single values per sample. Comparisons of source proportions across facilities were performed using Wilcoxon rank-sum tests with Benjamini-Hochberg correction.

## List of Abbreviations

ASV: Amplicon Sequence Variant; high-resolution DNA sequences used to identify microbial taxa.
BSA: Bovine Serum Albumin; a protein added to PCR reactions to enhance performance.
FEAST: Fast Expectation-Maximization for Microbial Source Tracking; a tool for estimating microbial source contributions.
PCoA: Principal Coordinate Analysis; a method for visualizing differences in microbial communities.
PCR: Polymerase Chain Reaction; a technique used to amplify DNA.
PERMANOVA: Permutational Multivariate Analysis of Variance; a statistical test for group differences based on distance metrics.
RAS: Recirculating Aquaculture System; a closed-loop water filtration system used in aquaculture.
SIMPER: Similarity Percentage Analysis; a method to determine which taxa contribute most to differences between groups.

## Declarations

### Ethics approval and consent to participate

Zebrafish at the Ore1 and Ore2 facilities were sampled and euthanized with approval from the University of Oregon’s Institutional Animal Care and Use Committee under protocol AUP-20-16. Zebrafish at the Norwegian facilities were euthanized in accordance with Norwegian regulations for animal research under approval from the Norwegian Food Safety Authority (Mattilsynet), permit #24125 (“Euthanasia of Zebrafish”).

### Consent for publication

Not applicable. This study does not contain any individual person’s data.

### Availability of data and materials

The datasets generated and/or analysed during the current study are available in the [NAME] repository, [PERSISTENT WEB LINK TO DATASETS].

### Authors’ contributions

KCE and BJMB conceived of the study. IB coordinated sample collection in Norway. KCE collected and processed samples and performed bioinformatic and statistical analyses. KCE wrote the manuscript and BJMB and IB provided critical revision and review. All authors read and approved the final manuscript.

## Acknowledgements

We thank the University of Oregon Zebrafish Facility, the Zebrafish International Resource Center, and the Jutfelt and Yaksi labs at the Norwegian University of Science and Technology for their valuable insights into facility operations and for granting access to sample collections. We are also grateful to Karen Adair for her assistance with sample processing and to Andrew H. Morris for support with data analysis. Additional thanks to the UO META Center, members of the Bohannan lab, and the IPAMA group for their thoughtful feedback and critical review of this research. Additionally, this material is based upon work supported by the National Science Foundation Graduate Research Fellowship under Grant No. 2236419.

## References

1. Teame T, Zhang Z, Ran C, Zhang H, Yang Y, Ding Q, et al. The use of zebrafish (Danio rerio) as biomedical models. Animal Frontiers. 2019;9:68–77. 10.1093/af/vfz020.

2. Wong S, Stephens WZ, Burns AR, Stagaman K, David LA, Bohannan BJM, et al. Ontogenetic Differences in Dietary Fat Influence Microbiota Assembly in the Zebrafish Gut. mBio. 2015;6:10.1128/mbio.00687-15. 10.1128/mbio.00687-15.

3. Adair KL, Douglas AE. Making a microbiome: the many determinants of host-associated microbial community composition. Current Opinion in Microbiology. 2017;35:23–9. 10.1016/j.mib.2016.11.002.

4. Kostic AD, Howitt MR, Garrett WS. Exploring host–microbiota interactions in animal models and humans. Genes Dev. 2013;27:701–18. 10.1101/gad.212522.112.

5. Kers JG, Velkers FC, Fischer EAJ, Hermes GDA, Lamot DM, Stegeman JA, et al. Take care of the environment: housing conditions affect the interplay of nutritional interventions and intestinal microbiota in broiler chickens. Animal Microbiome. 2019;1:10. 10.1186/s42523-019-0009-z.

6. Moraitou M, Forsythe A, Fellows Yates JA, Brealey JC, Warinner C, Guschanski K. Ecology, Not Host Phylogeny, Shapes the Oral Microbiome in Closely Related Species. Molecular Biology and Evolution. 2022;39:msac263. 10.1093/molbev/msac263.

7. Shigeno Y, Liu H, Sano C, Inoue R, Niimi K, Nagaoka K. Individual variations and effects of birth facilities on the fecal microbiome of laboratory-bred marmosets (Callithrix jacchus) assessed by a longitudinal study. PLOS ONE. 2022;17:e0273702. 10.1371/journal.pone.0273702.

8. Eisen JS. Chapter 1 - History of Zebrafish Research. In: Cartner SC, Eisen JS, Farmer SC, Guillemin KJ, Kent ML, Sanders GE, editors. The Zebrafish in Biomedical Research. Academic Press; 2020. p. 3–14. 10.1016/B978-0-12-812431-4.00001-4.

9. Stagaman K, Sharpton TJ, Guillemin K. Zebrafish microbiome studies make waves. Lab Anim. 2020;49:201–7. 10.1038/s41684-020-0573-6.

10. Parker KD, Albeke SE, Gigley JP, Goldstein AM, Ward NL. Microbiome Composition in Both Wild-Type and Disease Model Mice Is Heavily Influenced by Mouse Facility. Frontiers in Microbiology. 2018;9.

11. Rausch P, Basic M, Batra A, Bischoff SC, Blaut M, Clavel T, et al. Analysis of factors contributing to variation in the C57BL/6J fecal microbiota across German animal facilities. Int J Med Microbiol. 2016;306:343–55. 10.1016/j.ijmm.2016.03.004.

12. Flynn JK, Ortiz AM, Herbert R, Brenchley JM. Host Genetics and Environment Shape the Composition of the Gastrointestinal Microbiome in Nonhuman Primates. Microbiology Spectrum. 2022;11:e02139–22. 10.1128/spectrum.02139-22.

13. Breen P, Winters AD, Nag D, Ahmad MM, Theis KR, Withey JH. Internal Versus External Pressures: Effect of Housing Systems on the Zebrafish Microbiome. Zebrafish. 2019;16:388– 400. 10.1089/zeb.2018.1711.

14. Roeselers G, Mittge EK, Stephens WZ, Parichy DM, Cavanaugh CM, Guillemin K, et al. Evidence for a core gut microbiota in the zebrafish. ISME J. 2011;5:1595–608. 10.1038/ismej.2011.38.

15. Sharpton TJ, Stagaman K, Sieler MJ, Arnold HK, Davis EW. Phylogenetic Integration Reveals the Zebrafish Core Microbiome and Its Sensitivity to Environmental Exposures. Toxics. 2021;9:10. 10.3390/toxics9010010.

16. Attramadal KJK, Truong TMH, Bakke I, Skjermo J, Olsen Y, Vadstein O. RAS and microbial maturation as tools for K-selection of microbial communities improve survival in cod larvae. Aquaculture. 2014;432:483–90. 10.1016/j.aquaculture.2014.05.052.

17. Attramadal KJK, Minniti G, Øie G, Kjørsvik E, Østensen M-A, Bakke I, et al. Microbial maturation of intake water at different carrying capacities affects microbial control in rearing tanks for marine fish larvae. Aquaculture. 2016;457:68–72. 10.1016/j.aquaculture.2016.02.015.

18. Salvesen I, Skjermo J, Vadstein O. Growth of turbot (Scophthalmus maximus L.) during first feeding in relation to the proportion of r/K-strategists in the bacterial community of the rearing water. Aquaculture. 1999;175:337–50. 10.1016/S0044-8486(99)00110-6.

19. Clarke KR. Non-parametric multivariate analyses of changes in community structure. Australian Journal of Ecology. 1993;18:117–43. 10.1111/j.1442-9993.1993.tb00438.x.

20. Shenhav L, Thompson M, Joseph TA, Briscoe L, Furman O, Bogumil D, et al. FEAST: fast expectation-maximization for microbial source tracking. Nat Methods. 2019;16:627–32. 10.1038/s41592-019-0431-x.

21. Ofek T, Lalzar M, Laviad-Shitrit S, Izhaki I, Halpern M. Comparative Study of Intestinal Microbiota Composition of Six Edible Fish Species. Frontiers in Microbiology. 2021;12.

22. Ramírez C, Coronado J, Silva A, Romero J. Cetobacterium Is a Major Component of the Microbiome of Giant Amazonian Fish (Arapaima gigas) in Ecuador. Animals (Basel). 2018;8:189. 10.3390/ani8110189.

23. Yajima D, Fujita H, Hayashi I, Shima G, Suzuki K, Toju H. Core species and interactions prominent in fish-associated microbiome dynamics. Microbiome. 2023;11:53. 10.1186/s40168-023-01498-x.

24. Stephens WZ, Burns AR, Stagaman K, Wong S, Rawls JF, Guillemin K, et al. The composition of the zebrafish intestinal microbial community varies across development. ISME J. 2015;10:644–54. 10.1038/ismej.2015.140.

25. Gaulke CA, Martins ML, Watral VG, Humphreys IR, Spagnoli ST, Kent ML, et al. A longitudinal assessment of host-microbe-parasite interactions resolves the zebrafish gut microbiome’s link to Pseudocapillaria tomentosa infection and pathology. Microbiome. 2019;7:10. 10.1186/s40168-019-0622-9.

26. Wang A, Zhang Z, Ding Q, Yang Y, Bindelle J, Ran C, et al. Intestinal Cetobacterium and acetate modify glucose homeostasis via parasympathetic activation in zebrafish. Gut Microbes. 2020;13:1–15. 10.1080/19490976.2021.1900996.

27. Xie M, Hao Q, Xia R, Olsen RE, Ringø E, Yang Y, et al. Nuclease-Treated Stabilized Fermentation Product of Cetobacterium somerae Improves Growth, Non-specific Immunity, and Liver Health of Zebrafish (Danio rerio). Frontiers in Nutrition. 2022;9.

28. Quero GM, Piredda R, Basili M, Maricchiolo G, Mirto S, Manini E, et al. Host-associated and Environmental Microbiomes in an Open-Sea Mediterranean Gilthead Sea Bream Fish Farm. Microb Ecol. 2023;86:1319–30. 10.1007/s00248-022-02120-7.

30. Manchanayake T, Salleh A, Amal MNA, Yasin ISM, Zamri-Saad M. Pathology and pathogenesis of Vibrio infection in fish: A review. Aquaculture Reports. 2023;28:101459. 10.1016/j.aqrep.2022.101459.

31. Ringø E, Løvmo L, Kristiansen M, Bakken Y, Salinas I, Myklebust R, et al. Lactic acid bacteria vs. pathogens in the gastrointestinal tract of fish: a review. Aquaculture Research. 2010;41:451–67. 10.1111/j.1365-2109.2009.02339.x.

32. Sylvain F-É, Leroux N, Normandeau E, Custodio J, Mercier P-L, Bouslama S, et al. Fish-microbe systems in the hostile but highly biodiverse Amazonian blackwaters. 2022;:2022.10.22.513327. 10.1101/2022.10.22.513327.

33. Babenzien H-D, Cypionka H. Nevskia. In: Bergey’s Manual of Systematics of Archaea and Bacteria. John Wiley & Sons, Ltd; 2015. p. 1–6. 10.1002/9781118960608.gbm01233.

34. Wang Y, Fei S, Gao X, Wu H, Hong Z, Hu K. Mechanical abrasion stimulation: Altered epidermal mucus composition and microbial community in grass carp (*Ctenopharyngodon idella*). Aquaculture Reports. 2024;35:101936. 10.1016/j.aqrep.2024.101936.

35. François-Étienne S, Nicolas L, Eric N, Jaqueline C, Pierre-Luc M, Sidki B, et al. Important role of endogenous microbial symbionts of fish gills in the challenging but highly biodiverse Amazonian blackwaters. Nat Commun. 2023;14:3903. 10.1038/s41467-023-39461-x.

36. Zhao Y, Qin Z, Huang Z, Bao Z, Luo T, Jin Y. Effects of polyethylene microplastics on the microbiome and metabolism in larval zebrafish. Environmental Pollution. 2021;282:117039. 10.1016/j.envpol.2021.117039.

37. Bhat SV, Maughan H, Cameron ADS, Yost CK. Phylogenomic analysis of the genus Delftia reveals distinct major lineages with ecological specializations. Microbial Genomics. 2022;8. 10.1099/mgen.0.000864.

38. Goldschmidt-Clermont E, Wahli T, Frey J, Burr SE. Identification of bacteria from the normal flora of perch, Perca fluviatilis L., and evaluation of their inhibitory potential towards Aeromonas species. Journal of Fish Diseases. 2008;31:353–9. 10.1111/j.1365-2761.2008.00912.x.

39. Stressmann FA, Bernal-Bayard J, Perez-Pascual D, Audrain B, Rendueles O, Briolat V, et al. Mining zebrafish microbiota reveals key community-level resistance against fish pathogen infection. ISME J. 2021;15:702–19. 10.1038/s41396-020-00807-8.

40. Vargas O, Gutiérrez MS, Caruffo M, Valderrama B, Medina DA, García K, et al. Probiotic Yeasts and Vibrio anguillarum Infection Modify the Microbiome of Zebrafish Larvae. Frontiers in Microbiology. 2021;12.

41. Presta L, Bosi E, Fondi M, Maida I, Perrin E, Miceli E, et al. Phenotypic and genomic characterization of the antimicrobial producer *Rheinheimera* sp. EpRS3 isolated from the medicinal plant *Echinacea purpurea*: insights into its biotechnological relevance. Research in Microbiology. 2017;168:293–305. 10.1016/j.resmic.2016.11.001.

42. Sheu S-Y, Chen W-T, Young C-C, Chen W-M. Rheinheimera coerulea sp. nov., isolated from a freshwater creek, and emended description of genus Rheinheimera Brettar et al. 2002. International Journal of Systematic and Evolutionary Microbiology. 2018;68:2340–7. 10.1099/ijsem.0.002838.

43. Phelps D, Brinkman NE, Keely SP, Anneken EM, Catron TR, Betancourt D, et al. Microbial colonization is required for normal neurobehavioral development in zebrafish. Sci Rep. 2017;7:11244. 10.1038/s41598-017-10517-5.

44. Weitekamp CA, Phelps D, Swank A, McCord J, Sobus JR, Catron T, et al. Triclosan-Selected Host-Associated Microbiota Perform Xenobiotic Biotransformations in Larval Zebrafish. Toxicological Sciences. 2019;172:109–22. 10.1093/toxsci/kfz166.

45. Young GR, Sherry A, Smith DL. Built environment microbiomes transition from outdoor to human-associated communities after construction and commissioning. Sci Rep. 2023;13:15854. 10.1038/s41598-023-42427-0.

46. Meadow JF, Altrichter AE, Bateman AC, Stenson J, Brown GZ, Green JL, et al. Humans differ in their personal microbial cloud. PeerJ. 2015;3:e1258. 10.7717/peerj.1258.

47. D’Gama PP, Qiu T, Cosacak MI, Rayamajhi D, Konac A, Hansen JN, et al. Diversity and function of motile ciliated cell types within ependymal lineages of the zebrafish brain. Cell Rep. 2021;37:109775. 10.1016/j.celrep.2021.109775.

48. Lin L, Spoor MS, Gerth AJ, Brody SL, Peng SL. Modulation of Th1 Activation and Inflammation by the NF-κB Repressor Foxj1. Science. 2004;303:1017–20. 10.1126/science.1093889.

49. Vadstein O, Attramadal KJK, Bakke I, Olsen Y. K-Selection as Microbial Community Management Strategy: A Method for Improved Viability of Larvae in Aquaculture. Front Microbiol. 2018;9. 10.3389/fmicb.2018.02730.

50. Lu H, Li P, Huang X, Wang CH, Li M, Xu ZZ. Zebrafish model for human gut microbiome-related studies: advantages and limitations. Medicine in Microecology. 2021;8:100042. 10.1016/j.medmic.2021.100042.

51. Sprague J, Doerry E, Douglas S, Westerfield M. The Zebrafish Information Network (ZFIN): a resource for genetic, genomic and developmental research. Nucleic Acids Research. 2001;29:87–90. 10.1093/nar/29.1.87.

52. Trevarrow B, Robison B. Genetic Backgrounds, Standard Lines, and Husbandry of Zebrafish. In: Methods in Cell Biology. Academic Press; 2004. p. 599–616. 10.1016/S0091-679X(04)77032-6.

53. Blazewicz SJ, Barnard RL, Daly RA, Firestone MK. Evaluating rRNA as an indicator of microbial activity in environmental communities: limitations and uses. The ISME Journal. 2013;7:2061–8. 10.1038/ismej.2013.102.

54. Lebov JF, Schlomann BH, Robinson CD, Bohannan BJM. Phenotypic Parallelism during Experimental Adaptation of a Free-Living Bacterium to the Zebrafish Gut. mBio. 2020;11:10.1128/mbio.01519-20. 10.1128/mbio.01519-20.

55. Angert ER. Epulopiscium spp. Trends in Microbiology. 2022;30:97–8. 10.1016/j.tim.2021.11.004.

56. Krotman Y, Yergaliyev TM, Alexander Shani R, Avrahami Y, Szitenberg A. Dissecting the factors shaping fish skin microbiomes in a heterogeneous inland water system. Microbiome. 2020;8:9. 10.1186/s40168-020-0784-5.

57. Burns AR, Miller E, Agarwal M, Rolig AS, Milligan-Myhre K, Seredick S, et al. Interhost dispersal alters microbiome assembly and can overwhelm host innate immunity in an experimental zebrafish model. Proceedings of the National Academy of Sciences. 2017;114:11181–6. 10.1073/pnas.1702511114.

58. Stagaman K, Burns AR, Guillemin K, Bohannan BJ. The role of adaptive immunity as an ecological filter on the gut microbiota in zebrafish. ISME J. 2017;11:1630–9. 10.1038/ismej.2017.28.

59. Almeida AR, Alves M, Domingues I, Henriques I. The impact of antibiotic exposure in water and zebrafish gut microbiomes: A 16S rRNA gene-based metagenomic analysis. Ecotoxicol Environ Saf. 2019;186:109771. 10.1016/j.ecoenv.2019.109771.

60. Schmidt VT, Smith KF, Melvin DW, Amaral-Zettler LA. Community assembly of a euryhaline fish microbiome during salinity acclimation. Molecular Ecology. 2015;24:2537–50. 10.1111/mec.13177.

61. Bakke I, Skjermo J, Vo TA, Vadstein O. Live feed is not a major determinant of the microbiota associated with cod larvae (Gadus morhua). Environmental Microbiology Reports. 2013;5:537–48. 10.1111/1758-2229.12042.

62. Bik EM, Costello EK, Switzer AD, Callahan BJ, Holmes SP, Wells RS, et al. Marine mammals harbor unique microbiotas shaped by and yet distinct from the sea. Nat Commun. 2016;7:10516. 10.1038/ncomms10516.

63. Evens K. Fish Out of Water: Understanding the Impacts of Regional Species Pool Variation on Local Community Assembly in a Host-Microbiome Model System. 2024.

64. Mandal RK, Denny JE, Waide ML, Li Q, Bhutiani N, Anderson CD, et al. Temporospatial shifts within commercial laboratory mouse gut microbiota impact experimental reproducibility. BMC Biology. 2020;18:83. 10.1186/s12915-020-00810-7.

65. R Core Team. R: A Language and Environment for Statistical Computing. 2024.

66. Callahan BJ, McMurdie PJ, Rosen MJ, Han AW, Johnson AJA, Holmes SP. DADA2: High-resolution sample inference from Illumina amplicon data. Nat Methods. 2016;13:581–3. 10.1038/nmeth.3869.

67. Quast C, Pruesse E, Yilmaz P, Gerken J, Schweer T, Yarza P, et al. The SILVA ribosomal RNA gene database project: improved data processing and web-based tools. Nucleic Acids Research. 2012;41:D590–6. 10.1093/nar/gks1219.

68. Wang Q, Garrity GM, Tiedje JM, Cole JR. Naive Bayesian classifier for rapid assignment of rRNA sequences into the new bacterial taxonomy. Appl Environ Microbiol. 2007;73:5261–7. 10.1128/AEM.00062-07.

69. Davis NM, Proctor DM, Holmes SP, Relman DA, Callahan BJ. Simple statistical identification and removal of contaminant sequences in marker-gene and metagenomics data. Microbiome. 2018;6:226. 10.1186/s40168-018-0605-2.

70. Bray JR, Curtis JT. An Ordination of the Upland Forest Communities of Southern Wisconsin. Ecological Monographs. 1957;27:325–49. 10.2307/1942268.

71. Oksanen J, Simpson GL, Blanchet FG, Kindt R, Legendre P, Minchin PR, et al. vegan: Community Ecology Package. 2025.

72. Wickham H, Chang W, Henry L, Pedersen TL, Takahashi K, Wilke C, et al. ggplot2: Create Elegant Data Visualisations Using the Grammar of Graphics. 2025.

73. Hollander M, Wolfe DA, Chicken E. Nonparametric Statistical Methods. John Wiley & Sons; 2013.

74. Benjamini Y, Hochberg Y. Controlling the False Discovery Rate: A Practical and Powerful Approach to Multiple Testing. Journal of the Royal Statistical Society: Series B (Methodological). 1995;57:289–300. 10.1111/j.2517-6161.1995.tb02031.x.

75. Wilcoxon F. Individual Comparisons by Ranking Methods. Biometrics Bulletin. 1945;1:80– 3. 10.2307/3001968.

76. Martinez Arbizu P. pmartinezarbizu/pairwiseAdonis. 2025.

77. Teunisse GM. Fantaxtic - Nested Bar Plots for Phyloseq Data. 2022.

