## Supplemental Tables 1, 3, 5 for "Aquaculture facility-specific microbiota shape the zebrafish gut microbiome"

### SUPPLEMENTARY FIGURES 1, 3, 5

**Supp. Table 1 –** *PERMANOVA results for water microbiota comparisons between zebrafish facilities. Statistical significance was assessed using Bray-Curtis dissimilarity and unweighted UniFrac distance metrics under both hierarchical and constrained models. Significant differences (p < 0.05) are indicated.*

Tank Water PERMANOVA Results (Bray-Curtis)

| | Df | SumOfSqs | R2 | F | Pr(>F) |
|---|---|---|---|---|---|
| **A) Hierarchical PERMANOVA: Location, Facility, and Genotype Effects** | | | | | |
| Location | 1 | 3.5379237 | 0.2044461 | 17.139118 | 0.001 |
| Geno_Status...2 | 1 | 0.7403404 | 0.0427821 | 3.586505 | 0.002 |
| Location:Facility | 3 | 3.9440028 | 0.2279122 | 6.368776 | 0.001 |
| Residual...4 | 44 | 9.0826518 | 0.5248595 | NA | NA |
| Total...5 | 49 | 17.3049188 | 1.0000000 | NA | NA |
| **B) Constrained PERMANOVA: Genotype Effect within Facilities** | | | | | |
| Geno_Status...6 | 1 | 2.7283727 | 0.1576646 | 8.984426 | 0.567 |
| Residual...7 | 48 | 14.5765460 | 0.8423354 | NA | NA |
| Total...8 | 49 | 17.3049188 | 1.0000000 | NA | NA |

Tank Water PERMANOVA Results (Unweighted UniFrac)

| | Df | SumOfSqs | R2 | F | Pr(>F) |
|---|---|---|---|---|---|
| **A) Hierarchical PERMANOVA: Location, Facility, and Genotype Effects** | | | | | |
| Location | 1 | 1.6746172 | 0.1286687 | 8.808158 | 0.001 |
| Geno_Status...2 | 1 | 0.6197889 | 0.0476213 | 3.259968 | 0.001 |
| Location:Facility | 3 | 2.3552145 | 0.1809622 | 4.129322 | 0.001 |
| Residual...4 | 44 | 8.3653315 | 0.6427478 | NA | NA |
| Total...5 | 49 | 13.0149521 | 1.0000000 | NA | NA |
| **B) Constrained PERMANOVA: Genotype Effect within Facilities** | | | | | |
| Geno_Status...6 | 1 | 1.5945924 | 0.1225200 | 6.702104 | 0.427 |
| Residual...7 | 48 | 11.4203597 | 0.8774800 | NA | NA |
| Total...8 | 49 | 13.0149521 | 1.0000000 | NA | NA |

**Supp. Table 3 –** *PERMANOVA results for fish microbiota comparisons between zebrafish facilities. Statistical significance was assessed using Bray-Curtis dissimilarity and unweighted UniFrac distance metrics under both hierarchical and constrained models. Significant differences (p < 0.05) are indicated.*

Fish Gut PERMANOVA Results (Bray-Curtis)

| | Df | SumOfSqs | R2 | F | Pr(>F) |
|---|---|---|---|---|---|
| **A) Hierarchical PERMANOVA: Location, Facility, and Genotype Effects** | | | | | |
| Location | 1 | 9.791717 | 0.2190837 | 44.485881 | 0.001 |
| Geno_Status...2 | 1 | 1.561720 | 0.0349425 | 7.095229 | 0.001 |
| Location:Facility | 3 | 5.166642 | 0.1156005 | 7.824389 | 0.001 |
| Residual...4 | 128 | 28.173877 | 0.6303733 | NA | NA |
| Total...5 | 133 | 44.693955 | 1.0000000 | NA | NA |
| **B) Constrained PERMANOVA: Genotype Effect within Facilities** | | | | | |
| Geno_Status...6 | 1 | 5.581787 | 0.1248891 | 18.838022 | 0.167 |
| Residual...7 | 132 | 39.112168 | 0.8751109 | NA | NA |
| Total...8 | 133 | 44.693955 | 1.0000000 | NA | NA |

Fish Gut PERMANOVA Results (Unweighted UniFrac)

| | Df | SumOfSqs | R2 | F | Pr(>F) |
|---|---|---|---|---|---|
| **A) Hierarchical PERMANOVA: Location, Facility, and Genotype Effects** | | | | | |
| Location | 1 | 5.002274 | 0.1235252 | 21.752720 | 0.001 |
| Geno_Status...2 | 1 | 1.920098 | 0.0474145 | 8.349673 | 0.001 |
| Location:Facility | 3 | 4.138626 | 0.1021984 | 5.999031 | 0.001 |
| Residual...4 | 128 | 29.434985 | 0.7268619 | NA | NA |
| Total...5 | 133 | 40.495983 | 1.0000000 | NA | NA |
| **B) Constrained PERMANOVA: Genotype Effect within Facilities** | | | | | |
| Geno_Status...6 | 1 | 4.222407 | 0.1042673 | 15.365393 | 0.742 |
| Residual...7 | 132 | 36.273576 | 0.8957327 | NA | NA |
| Total...8 | 133 | 40.495983 | 1.0000000 | NA | NA |

**Supp. Figure 5 -** *Bray-Curtis dissimilarity boxplots of paired fish and water samples separated by facility*: Tukey-style box and whisker plot featuring pairwise beta-diversity values fusing Bray-Curtis distance for fish gut microbiome samples separated by facility paired with 1) fish gut microbiome samples from the same facility, 2) Water microbiome samples from the same tank ('same tank'), 3) water from tanks within the same facility but not in the same tank ('same facility'), and 4) water from tanks in other facilities ('different facility'). Significance calculated with Kruskal-Wallis and pairwise Wilcoxon rank-sum test with Benjamini-Hochberg p-value correction (* = p < 0.05, ** = p < 0.01, *** = p < 0.001, **** = p < 0.0001, ns = non-significant).

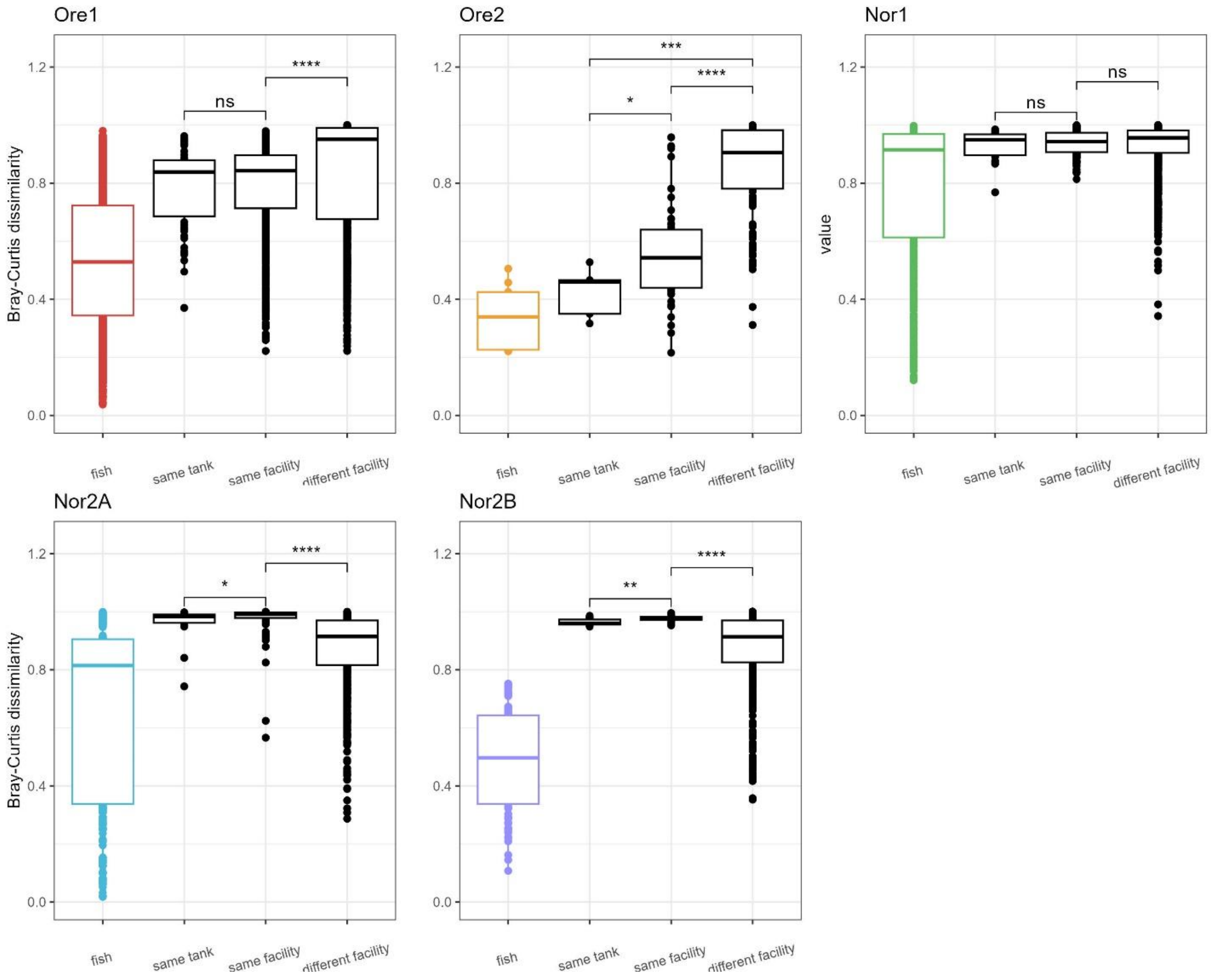
