## Supplemental Table 2 for "Aquaculture facility-specific microbiota shape the zebrafish gut microbiome"

Supplementary Table 2: Top 10 most significant ASVs contributing to water microbiome dissimilarity between facility comparisons (SIMPER analysis, $p < 0.05$)

| Facility Comparison | Genus | Average Contribution (%) | p-value | Rank |
|---|---|---|---|---|
| **Ore1 vs Ore2** | | | | |
| **Ore1 vs Ore2** | Cetobacterium | 13.78 | 0.011 | 1 |
| **Ore1 vs Ore2** | Vibrio | 4.85 | 0.001 | 4 |
| **Ore1 vs Ore2** | Plesiomonas | 2.58 | 0.001 | 8 |
| **Ore1 vs Ore2** | Chitinibacter | 1.33 | 0.001 | 10 |
| **Ore1 vs Ore2** | Chitinimonas | 0.97 | 0.001 | 11 |
| **Ore1 vs Ore2** | Crenobacter | 0.58 | 0.001 | 18 |
| **Ore1 vs Ore2** | Epulopiscium | 0.55 | 0.001 | 19 |
| **Ore1 vs Ore2** | Haliangium | 0.44 | 0.001 | 21 |
| **Ore1 vs Ore2** | Rhizorhapis | 0.34 | 0.001 | 24 |
| **Ore1 vs Ore2** | Romboutsia | 0.32 | 0.001 | 26 |
| **Ore1 vs Nor1** | | | | |
| **Ore1 vs Nor1** | Pseudomonas | 17.82 | 0.001 | 1 |
| **Ore1 vs Nor1** | Stenotrophomonas | 2.28 | 0.026 | 10 |
| **Ore1 vs Nor1** | Massilia | 1.79 | 0.001 | 11 |
| **Ore1 vs Nor1** | Variovorax | 0.99 | 0.001 | 16 |
| **Ore1 vs Nor1** | Nubsella | 0.89 | 0.010 | 17 |
| **Ore1 vs Nor1** | Deinococcus | 0.65 | 0.001 | 21 |
| **Ore1 vs Nor1** | Ottowia | 0.62 | 0.001 | 22 |
| **Ore1 vs Nor1** | Salinirepens | 0.61 | 0.040 | 23 |
| **Ore1 vs Nor1** | Alkanibacter | 0.51 | 0.001 | 26 |
| **Ore1 vs Nor1** | Sphingobacterium | 0.45 | 0.001 | 27 |
| **Ore1 vs Nor2A** | | | | |
| **Ore1 vs Nor2A** | Aeromonas | 6.00 | 0.027 | 3 |
| **Ore1 vs Nor2A** | Acidovorax | 5.72 | 0.001 | 4 |
| **Ore1 vs Nor2A** | Nevskia | 5.04 | 0.006 | 5 |
| **Ore1 vs Nor2A** | Limnobacter | 4.28 | 0.001 | 6 |
| **Ore1 vs Nor2A** | Luteimonas | 3.10 | 0.001 | 12 |
| **Ore1 vs Nor2A** | Stenotrophomonas | 2.29 | 0.033 | 15 |
| **Ore1 vs Nor2A** | Pelomonas | 2.13 | 0.004 | 16 |
| **Ore1 vs Nor2A** | Brevundimonas | 2.12 | 0.001 | 17 |
| **Ore1 vs Nor2A** | Rhodoferax | 1.21 | 0.001 | 21 |
| **Ore1 vs Nor2A** | Methyloversatilis | 1.19 | 0.003 | 22 |
| **Ore1 vs Nor2B** | | | | |

| Facility Comparison | Genus | Average Contribution (%) | p-value | Rank |
|---|---|---|---|---|
| **Ore1 vs Nor2B** | Rheinheimera | 16.91 | 0.001 | 1 |
| **Ore1 vs Nor2B** | Acinetobacter | 9.13 | 0.007 | 4 |
| **Ore1 vs Nor2B** | Fluviicola | 5.73 | 0.001 | 5 |
| **Ore1 vs Nor2B** | Polynucleobacter | 1.38 | 0.001 | 12 |
| **Ore1 vs Nor2B** | Sediminibacterium | 1.26 | 0.001 | 13 |
| **Ore1 vs Nor2B** | hgcI clade | 1.15 | 0.001 | 14 |
| **Ore1 vs Nor2B** | Candidatus Nitrosotenuis | 0.97 | 0.001 | 15 |
| **Ore1 vs Nor2B** | Bdellovibrio | 0.80 | 0.001 | 19 |
| **Ore1 vs Nor2B** | Candidatus Omnitrophus | 0.33 | 0.002 | 28 |
| **Ore1 vs Nor2B** | Cellvibrio | 0.24 | 0.001 | 34 |
| **Ore2 vs Nor1** | | | | |
| **Ore2 vs Nor1** | Cetobacterium | 18.11 | 0.001 | 1 |
| **Ore2 vs Nor1** | Pseudomonas | 18.10 | 0.001 | 2 |
| **Ore2 vs Nor1** | Vibrio | 4.52 | 0.007 | 3 |
| **Ore2 vs Nor1** | Plesiomonas | 2.73 | 0.003 | 5 |
| **Ore2 vs Nor1** | Stenotrophomonas | 2.29 | 0.033 | 8 |
| **Ore2 vs Nor1** | Massilia | 1.80 | 0.001 | 9 |
| **Ore2 vs Nor1** | Variovorax | 1.00 | 0.001 | 15 |
| **Ore2 vs Nor1** | Nubsella | 0.89 | 0.013 | 18 |
| **Ore2 vs Nor1** | Deinococcus | 0.68 | 0.001 | 19 |
| **Ore2 vs Nor1** | Ottowia | 0.64 | 0.001 | 21 |
| **Ore2 vs Nor2A** | | | | |
| **Ore2 vs Nor2A** | Cetobacterium | 20.33 | 0.001 | 1 |
| **Ore2 vs Nor2A** | Acidovorax | 5.71 | 0.001 | 2 |
| **Ore2 vs Nor2A** | Nevskia | 5.06 | 0.010 | 3 |
| **Ore2 vs Nor2A** | Vibrio | 4.70 | 0.007 | 4 |
| **Ore2 vs Nor2A** | Limnobacter | 4.57 | 0.001 | 5 |
| **Ore2 vs Nor2A** | Plesiomonas | 3.24 | 0.001 | 9 |
| **Ore2 vs Nor2A** | Luteimonas | 3.15 | 0.001 | 11 |
| **Ore2 vs Nor2A** | Stenotrophomonas | 2.29 | 0.048 | 13 |
| **Ore2 vs Nor2A** | Brevundimonas | 2.16 | 0.001 | 14 |
| **Ore2 vs Nor2A** | Pelomonas | 1.86 | 0.021 | 15 |

| Facility Comparison | Genus | Average Contribution (%) | p-value | Rank |
|---|---|---|---|---|
| **Ore2 vs Nor2B** | | | | |
| **Ore2 vs Nor2B** | Cetobacterium | 20.21 | 0.003 | 1 |
| **Ore2 vs Nor2B** | Rheinheimera | 17.06 | 0.001 | 2 |
| **Ore2 vs Nor2B** | Acinetobacter | 8.91 | 0.006 | 3 |
| **Ore2 vs Nor2B** | Fluviicola | 5.72 | 0.001 | 4 |
| **Ore2 vs Nor2B** | Vibrio | 5.20 | 0.034 | 5 |
| **Ore2 vs Nor2B** | Plesiomonas | 3.23 | 0.014 | 6 |
| **Ore2 vs Nor2B** | Polynucleobacter | 1.39 | 0.001 | 12 |
| **Ore2 vs Nor2B** | Sediminibacterium | 1.29 | 0.001 | 14 |
| **Ore2 vs Nor2B** | hgcI clade | 1.15 | 0.002 | 15 |
| **Ore2 vs Nor2B** | Candidatus Nitrosotenuis | 0.96 | 0.002 | 17 |
| **Nor2A vs Nor1** | | | | |
| **Nor2A vs Nor1** | Pseudomonas | 14.60 | 0.001 | 1 |
| **Nor2A vs Nor1** | Acidovorax | 5.38 | 0.001 | 2 |
| **Nor2A vs Nor1** | Nevskia | 4.91 | 0.019 | 3 |
| **Nor2A vs Nor1** | Delftia | 4.85 | 0.002 | 4 |
| **Nor2A vs Nor1** | Limnobacter | 4.31 | 0.001 | 5 |
| **Nor2A vs Nor1** | Luteimonas | 2.82 | 0.001 | 8 |
| **Nor2A vs Nor1** | Stenotrophomonas | 2.79 | 0.014 | 9 |
| **Nor2A vs Nor1** | Brevundimonas | 2.02 | 0.001 | 11 |
| **Nor2A vs Nor1** | Pelomonas | 1.86 | 0.035 | 13 |
| **Nor2A vs Nor1** | Massilia | 1.75 | 0.001 | 14 |
| **Nor2B vs Nor1** | | | | |
| **Nor2B vs Nor1** | Pseudomonas | 17.49 | 0.001 | 1 |
| **Nor2B vs Nor1** | Rheinheimera | 16.29 | 0.001 | 2 |
| **Nor2B vs Nor1** | Acinetobacter | 7.38 | 0.010 | 3 |
| **Nor2B vs Nor1** | Fluviicola | 5.52 | 0.001 | 4 |
| **Nor2B vs Nor1** | Massilia | 1.75 | 0.013 | 11 |

| Facility Comparison | Genus | Average Contribution (%) | p-value | Rank |
|---|---|---|---|---|
| **Nor2B vs Nor1** | Polynucleobacter | 1.33 | 0.017 | 14 |
| **Nor2B vs Nor1** | Sediminibacterium | 1.18 | 0.001 | 15 |
| **Nor2B vs Nor1** | hgcI clade | 1.10 | 0.005 | 18 |
| **Nor2B vs Nor1** | Variovorax | 0.98 | 0.048 | 19 |
| **Nor2B vs Nor1** | Candidatus Nitrosotenuis | 0.94 | 0.002 | 20 |
| **Nor2A vs Nor2B** | | | | |
| **Nor2A vs Nor2B** | Rheinheimera | 13.42 | 0.001 | 1 |
| **Nor2A vs Nor2B** | Acinetobacter | 8.32 | 0.003 | 2 |
| **Nor2A vs Nor2B** | Fluviicola | 5.61 | 0.001 | 3 |
| **Nor2A vs Nor2B** | Luteimonas | 3.06 | 0.006 | 8 |
| **Nor2A vs Nor2B** | Brevundimonas | 2.04 | 0.050 | 13 |
| **Nor2A vs Nor2B** | Polynucleobacter | 1.35 | 0.017 | 17 |
| **Nor2A vs Nor2B** | hgcI clade | 1.12 | 0.001 | 20 |
| **Nor2A vs Nor2B** | Sediminibacterium | 1.07 | 0.002 | 21 |
| **Nor2A vs Nor2B** | Candidatus Nitrosotenuis | 0.96 | 0.001 | 22 |
| **Nor2A vs Nor2B** | Bdellovibrio | 0.83 | 0.001 | 24 |
