## Supplemental Table 4 for "Aquaculture facility-specific microbiota shape the zebrafish gut microbiome"

Supplementary Table 3: Top 10 most significant ASVs contributing to fish gut microbiome dissimilarity between facility comparisons (SIMPER analysis, $p < 0.05$)

| Facility Comparison | Genus | Average Contribution (%) | p-value | Rank |
|---|---|---|---|---|
| **Ore1 vs Ore2** | | | | |
| **Ore1 vs Ore2** | Streptococcus | 5.94 | 0.001 | 3 |
| **Ore1 vs Ore2** | Enterococcus | 5.53 | 0.001 | 4 |
| **Ore1 vs Ore2** | Vagococcus | 0.74 | 0.008 | 12 |
| **Ore1 vs Ore2** | Epulopiscium | 0.33 | 0.016 | 19 |
| **Ore1 vs Ore2** | Gemmobacter | 0.10 | 0.003 | 41 |
| **Ore1 vs Ore2** | Clostridium sensu stricto 13 | 0.09 | 0.008 | 42 |
| **Ore1 vs Ore2** | Isosphaera | 0.09 | 0.012 | 43 |
| **Ore1 vs Ore2** | Rhodopirellula | 0.04 | 0.030 | 53 |
| **Ore1 vs Ore2** | Paeniglutamicibacter | 0.02 | 0.003 | 78 |
| **Ore1 vs Ore2** | Gaiella | 0.01 | 0.002 | 101 |
| **Ore1 vs Nor1** | | | | |
| **Ore1 vs Nor1** | Cetobacterium | 26.59 | 0.001 | 1 |
| **Ore1 vs Nor1** | Pseudomonas | 9.14 | 0.001 | 3 |
| **Ore1 vs Nor1** | Lactococcus | 3.47 | 0.001 | 5 |
| **Ore1 vs Nor1** | Streptococcus | 2.45 | 0.009 | 7 |
| **Ore1 vs Nor1** | Achromobacter | 2.38 | 0.001 | 8 |
| **Ore1 vs Nor1** | Pediococcus | 2.25 | 0.001 | 9 |
| **Ore1 vs Nor1** | Staphylococcus | 1.97 | 0.003 | 10 |
| **Ore1 vs Nor1** | Leuconostoc | 1.72 | 0.001 | 13 |
| **Ore1 vs Nor1** | Latilactobacillus | 1.23 | 0.008 | 14 |
| **Ore1 vs Nor1** | Nocardia | 1.10 | 0.008 | 15 |
| **Ore1 vs Nor2A** | | | | |
| **Ore1 vs Nor2A** | Vibrio | 23.29 | 0.001 | 1 |
| **Ore1 vs Nor2A** | Cetobacterium | 22.95 | 0.001 | 2 |
| **Ore1 vs Nor2A** | Bordetella | 1.25 | 0.041 | 8 |
| **Ore1 vs Nor2A** | Pseudonocardia | 0.48 | 0.035 | 18 |
| **Ore1 vs Nor2A** | Paucilactobacillus | 0.13 | 0.048 | 38 |
| **Ore1 vs Nor2A** | Gallicola | 0.08 | 0.043 | 46 |
| **Ore1 vs Nor2A** | Pedomicrobium | 0.05 | 0.020 | 53 |
| **Ore1 vs Nor2A** | Micrococcus | 0.00 | 0.005 | 164 |
| **Ore1 vs Nor2B** | | | | |
| **Ore1 vs Nor2B** | Aeromonas | 27.07 | 0.001 | 1 |
| **Ore1 vs Nor2B** | Cetobacterium | 26.43 | 0.001 | 2 |

| Facility Comparison | Genus | Average Contribution (%) | p-value | Rank |
|---|---|---|---|---|
| **Ore1 vs Nor2B** | Bacillus | 2.54 | 0.018 | 5 |
| **Ore1 vs Nor2B** | Crenobacter | 0.64 | 0.001 | 14 |
| **Ore1 vs Nor2B** | Chelativorans | 0.60 | 0.001 | 15 |
| **Ore1 vs Nor2B** | Marmoricola | 0.37 | 0.001 | 20 |
| **Ore1 vs Nor2B** | Pir4 lineage | 0.30 | 0.029 | 24 |
| **Ore1 vs Nor2B** | Chitinilyticum | 0.24 | 0.001 | 27 |
| **Ore1 vs Nor2B** | Hyphomicrobium | 0.21 | 0.045 | 30 |
| **Ore1 vs Nor2B** | Flavobacterium | 0.19 | 0.001 | 33 |
| **Ore2 vs Nor1** | | | | |
| **Ore2 vs Nor1** | Cetobacterium | 22.38 | 0.042 | 1 |
| **Ore2 vs Nor1** | Streptococcus | 5.90 | 0.003 | 5 |
| **Ore2 vs Nor1** | Enterococcus | 5.25 | 0.001 | 6 |
| **Ore2 vs Nor1** | Vagococcus | 0.87 | 0.001 | 16 |
| **Ore2 vs Nor1** | Epulopiscium | 0.27 | 0.050 | 27 |
| **Ore2 vs Nor1** | Gemmobacter | 0.09 | 0.006 | 41 |
| **Ore2 vs Nor1** | Clostridium sensu stricto 13 | 0.09 | 0.008 | 43 |
| **Ore2 vs Nor1** | Isosphaera | 0.08 | 0.026 | 45 |
| **Ore2 vs Nor1** | Paeniglutamicibacter | 0.02 | 0.040 | 83 |
| **Ore2 vs Nor1** | Clostridium sensu stricto 18 | 0.01 | 0.040 | 101 |
| **Ore2 vs Nor2A** | | | | |
| **Ore2 vs Nor2A** | Vibrio | 20.24 | 0.018 | 1 |
| **Ore2 vs Nor2A** | Streptococcus | 6.20 | 0.002 | 4 |
| **Ore2 vs Nor2A** | Enterococcus | 5.60 | 0.001 | 5 |
| **Ore2 vs Nor2A** | Vagococcus | 0.86 | 0.002 | 14 |
| **Ore2 vs Nor2A** | Gemmobacter | 0.09 | 0.013 | 23 |
| **Ore2 vs Nor2A** | Clostridium sensu stricto 13 | 0.09 | 0.013 | 24 |
| **Ore2 vs Nor2A** | Isosphaera | 0.08 | 0.026 | 25 |
| **Ore2 vs Nor2A** | Paeniglutamicibacter | 0.02 | 0.041 | 47 |
| **Ore2 vs Nor2A** | Clostridium sensu stricto 18 | 0.01 | 0.041 | 60 |
| **Ore2 vs Nor2A** | Desulfotomaculum | 0.01 | 0.041 | 64 |
| **Ore2 vs Nor2B** | | | | |

| Facility Comparison | Genus | Average Contribution (%) | p-value | Rank |
|---|---|---|---|---|
| **Ore2 vs Nor2B** | Aeromonas | 27.87 | 0.001 | 1 |
| **Ore2 vs Nor2B** | Streptococcus | 6.61 | 0.001 | 3 |
| **Ore2 vs Nor2B** | Enterococcus | 5.97 | 0.001 | 5 |
| **Ore2 vs Nor2B** | Vagococcus | 0.92 | 0.002 | 10 |
| **Ore2 vs Nor2B** | Crenobacter | 0.60 | 0.036 | 11 |
| **Ore2 vs Nor2B** | Chelativorans | 0.59 | 0.009 | 12 |
| **Ore2 vs Nor2B** | Marmoricola | 0.36 | 0.008 | 15 |
| **Ore2 vs Nor2B** | Epulopiscium | 0.29 | 0.048 | 17 |
| **Ore2 vs Nor2B** | Chitinilyticum | 0.24 | 0.029 | 18 |
| **Ore2 vs Nor2B** | Flavobacterium | 0.19 | 0.042 | 19 |
| **Nor2A vs Nor1** | | | | |
| **Nor2A vs Nor1** | Vibrio | 23.79 | 0.001 | 1 |
| **Nor2A vs Nor1** | Pseudomonas | 9.80 | 0.001 | 3 |
| **Nor2A vs Nor1** | Lactococcus | 3.22 | 0.022 | 5 |
| **Nor2A vs Nor1** | Stenotrophomonas | 2.75 | 0.045 | 6 |
| **Nor2A vs Nor1** | Delftia | 2.73 | 0.018 | 7 |
| **Nor2A vs Nor1** | Pediococcus | 2.18 | 0.005 | 10 |
| **Nor2A vs Nor1** | Leuconostoc | 1.67 | 0.006 | 13 |
| **Nor2A vs Nor1** | Bordetella | 1.26 | 0.033 | 14 |
| **Nor2A vs Nor1** | Limosilactobacillus | 0.27 | 0.005 | 28 |
| **Nor2A vs Nor1** | Pedomicrobium | 0.05 | 0.007 | 45 |
| **Nor2B vs Nor1** | | | | |
| **Nor2B vs Nor1** | Aeromonas | 20.17 | 0.001 | 1 |
| **Nor2B vs Nor1** | Pseudomonas | 9.41 | 0.014 | 2 |
| **Nor2B vs Nor1** | Lactococcus | 3.43 | 0.040 | 4 |
| **Nor2B vs Nor1** | Bacillus | 2.48 | 0.030 | 7 |
| **Nor2B vs Nor1** | Pediococcus | 2.32 | 0.014 | 10 |
| **Nor2B vs Nor1** | Leuconostoc | 1.78 | 0.017 | 14 |

| Facility Comparison | Genus | Average Contribution (%) | p-value | Rank |
|---|---|---|---|---|
| **Nor2B vs Nor1** | Candidatus Nitrocosmicus | 1.50 | 0.016 | 15 |
| **Nor2B vs Nor1** | Mycobacterium | 0.78 | 0.011 | 16 |
| **Nor2B vs Nor1** | Crenobacter | 0.67 | 0.001 | 17 |
| **Nor2B vs Nor1** | Chelativorans | 0.59 | 0.001 | 19 |
| **Nor2A vs Nor2B** | | | | |
| **Nor2A vs Nor2B** | Vibrio | 23.66 | 0.001 | 1 |
| **Nor2A vs Nor2B** | Aeromonas | 23.65 | 0.001 | 2 |
| **Nor2A vs Nor2B** | Bacillus | 3.34 | 0.002 | 5 |
| **Nor2A vs Nor2B** | Crenobacter | 0.62 | 0.001 | 12 |
| **Nor2A vs Nor2B** | Chelativorans | 0.58 | 0.001 | 13 |
| **Nor2A vs Nor2B** | Marmoricola | 0.36 | 0.001 | 18 |
| **Nor2A vs Nor2B** | Chitinilyticum | 0.23 | 0.001 | 21 |
| **Nor2A vs Nor2B** | Flavobacterium | 0.18 | 0.001 | 22 |
| **Nor2A vs Nor2B** | Pirellula | 0.17 | 0.001 | 23 |
| **Nor2A vs Nor2B** | Nannocystis | 0.14 | 0.002 | 26 |
