## Supplemental Table 6 for "Aquaculture facility-specific microbiota shape the zebrafish gut microbiome"

| | Facility | | | | | | |
|---|---|---|---|---|---|---|---|
| **Genotype** | **Ore1** | **Ore2** | **Nor1** | **Nor2A** | **Nor2B** | **WT/GM Classification** | **Notes** |
| AB | X | X | | | | WT | Standard laboratory strain, commonly used wild-type reference |
| ABC | X | | | | | WT | AB-related strain maintained without round robin breeding |
| WT | | | X | | X | WT | Wild-type control strain; no additional strain information |
| NACRE | | | | X | | GM | MITF gene pigmentation mutant, transparent/translucent for imaging |
| HucGcamp6 | | | X | X | | GM | Neuronal calcium indicator line (HuC promoter drives GCaMP6) |
| GMNC1 x GMNC | | | X | | | GM | Seizure/epilepsy model line (elevated photic response, network decay) |
| OMP x ChR2/3/4 | | | X | | | GM | Olfactory neuron optogenetics lines (olfactory marker protein promoter) |
| R2 | | | X | | | GM | Facility-specific line designation |
| Elipsa | | | X | | | GM | Loss-of-function mutation of traf3ip1 |
| npygRNA2 x vas x Gcamp6 | | | X | | | GM | Seizure/epilepsy model line (elevated photic response, network decay) |
| vas x Gcamp6 | | | X | | | GM | Germline calcium indicator (vasa promoter drives GCaMP6) |
| Foxjlb x Fauf x vas x GCatiP6s | | | X | | | GM | Multi-transgenic line for cilia/germline imaging |
| 6S | | | X | | | GM | Facility-specific line designation |
